# Synergistic activation of the human phosphate exporter XPR1 by KIDINS220 and inositol pyrophosphate

**DOI:** 10.1101/2024.12.22.630011

**Authors:** Peng Zuo, Weize Wang, Zonglin Dai, Jiye Zheng, Shang Yu, Guangxi Wang, Yue Yin, Ling Liang, Yuxin Yin

**Affiliations:** Institute of Systems Biomedicine, Department of Pathology, Beijing Key Laboratory of Tumor Systems Biology, School of Basic Medical Sciences, Peking University Health Science Center, Beijing 100191, China; Peking-Tsinghua Center for Life Sciences, Peking University, Beijing, 100871, China; Department of Biophysics, School of Basic Medical Sciences, Peking University Health Science Center, Beijing 100191, China; Department of Pharmacology, School of Basic Medical Sciences, Peking University Health Science Center, Beijing 100191, China; Institute of Precision Medicine, Peking University Shenzhen Hospital, Shenzhen 518036, China

## Abstract

Inorganic phosphate (Pi) is fundamental to life, and its intracellular concentration must be tightly regulated to prevent toxicity. XPR1, the only known phosphate exporter, plays a crucial role in maintaining this delicate balance. However, the mechanisms underlying the function and regulation of XPR1 remain elusive until now. Here we present cryo-electron microscopy structures of the human XPR1-KIDINS220 complex in both substrate-free closed states and substrate-bound outward-open states, as well as the structure of an XPR1 mutant alone in a substrate-bound inward-facing state. In the presence of inositol hexaphosphate (InsP6) and phosphate, the XPR1-KIDINS220 complex adopts an outward-open conformation. InsP6 binds both the SPX domain and the peripheral juxtamembrane regions of XPR1, indicative of an active phosphate-export state. Conversely, in the absence of either phosphate or InsP6, the complex assumes a closed state, where the extracellular half of transmembrane 9 occupies the outward cavity, and a C-terminal plug-in loop blocks the intracellular cavity. Notably, XPR1 without KIDINS220 adopts a closed state despite the presence of phosphate and InsP6. The functional mutagenesis experiments further demonstrate that InsP6, whose concentrations fluctuate in response to Pi availability, functions synergistically with KIDINS220 to regulate the phosphate export activity of XPR1. These findings not only elucidate the intricate mechanisms of cellular phosphate regulation but also hold promise for the development of targeted therapies for ovarian cancer, where XPR1 plays a significant role.

## Introduction

Phosphorus (P) is an essential element for life. It is widely involved in the composition of important biological macromolecules like DNA and phospholipids, as well as various cell biological processes including energy delivery and signaling transduction^1^. Inorganic phosphate (Pi, H_2_PO_4_^−^ or HPO_4_^2−^) is the primary form of phosphorus absorption for all living organisms, thus acquiring sufficient but not excessive Pi is crucial for cell metabolism and survival^2^. To maintain phosphate homeostasis, bacteria, fungi, plants, and animals have developed sophisticated regulatory networks to detect and manage intracellular phosphate levels. Among these mechanisms, multiple plasma membrane-localized Pi transporters play crucial roles as key functional executors.^2–5^. In vertebrates, two types of sodium-phosphate cotransporters are essential for cellular phosphate uptake and exhibit tissue-specific distribution: PiT1 and PiT2, encoded by the *SLC20A1* and *SLC20A2* genes, as well as NaPi-IIa, NaPi-IIb, and NaPi-IIc, encoded by the *SLC34A1-3* genes^6,7^, respectively. Regarding phosphate export, the xenotropic and polytropic retrovirus receptor 1 (XPR1, also known as SLC53A1) has been identified as the sole phosphate exporter in metazoans^8^. These transporters collectively contribute to the maintenance of intracellular phosphate homeostasis of vertebrates.

Human XPR1 was initially identified as the cell surface receptor for xenotropic and polytropic murine leukemia viruses^9–11^, similar to how PiT1 and PiT2 serve as receptors for Glvr and Ram retroviruses^12^. XPR1 has been reported to be widely expressed in mammalian cells, excluding erythrocytes, and it mediates phosphate export in vertebrates^8^, which was also confirmed in a heterologous expression system of wild tobacco^13^. The phosphate export function of XPR1 plays an important role in the survival of mammals, as *Xpr1*deletion causes lethality in mouse pups^14,15^. XPR1 has also been reported to participate in the physiological processes of various mammalian tissues, including renal tubules^14^, placenta^15^, platelet^16^, pancreatic β-cells^17^ and aorta^18^. Mutations of *XPR1* gene may cause several human diseases, such as Primary familial brain calcification (PFBC)^19,20^ and renal Fanconi syndrome^14,21^. Moreover, over-expression or mutations of *XPR1* gene have also been reported to be associated with the tumorigenisis of multiple cancers^22,23^. Therefore, investigation on the phosphate transport mechanism of XPR1 has substantial physiological and pathological significance.

Mammalian XPR1 has been characterized to share similarity in sequence and domain architecture with SYG1 in yeasts and PHO1 in plants. The analogous features include a N-terminal cytoplasmic SPX domain (named after SYG1, PHO81 and XPR1) and a C-terminal transmembrane EXS domain (named after Erd1, XPR1 and SYG1)^9–11,24,25^. While SYG1 is considered to be a putative Pi exporter^8^, PHO1 has been identified to mediate the Pi transfer from roots to shoots through the EXS domain^24–26^. Instead, the N-terminal SPX domain regulates the transport process through its binding with inositol polyphosphate^27^. And studies on human XPR1 mediated Pi export activity have illustrated that inositol pyrophosphate InsP8 serves as the direct regulatory molecule for XPR1’s SPX domain^28,29^. Moreover, it has been reported that the phosphate export function of XPR1 requires the participation of other proteins. KIDINS220, a neuronal substrate of protein Kinase D^30^, has been shown to be a co-factor for the phosphate export function of XPR1, and their complex could potentially serve as a viable therapeutic target in ovarian and uterine cancer^31^. Additionally, XPR1-mediated phosphate efflux has also been discovered to interplay with the phosphate uptake activities of PiT1 and PiT2 with the assistance of inositol pyrophosphates^32,33^. In light of the researches above, the roles of these regulatory molecules and protein partners in XPR1’s function and cellular phosphate homeostasis deserve further investigation.

Here, we resolved the cryo-electron microscopy (cryo-EM) structures of human XPR1 with or without the presence of InsP6 or KIDINS220 ankyrin-repeats, and revealed that only in the presence of both ligands and substrate KH_2_PO_4_, the conformation of XPR1 could transit from a closed state to an outward-open state. Besides, we explored the significance of the C-terminal plug-in loop (Glu622/Phe623 motif) of XPR1 in its phosphate export activity and resolved the structure of the XPR1-E622A/F623A mutant in the inward-facing state. Interestingly, we identified two InsP6 binding sites within XPR1, one at the interface between the SPX domains and the other between the SPX and the TMD. Moreover, we investigated the roles of several critical residues of XPR1 in the InsP6 binding pockets as well as in the phosphate transport pathway by utilizing ^[32^P^]^ KH_2_PO_4_ efflux assays. To sum up the results above, we proposed a “rocking-bundle”, alternating-access mechanism to depict the transport cycle of phosphate export mediated by XPR1 in addition with KIDINS220 and InsP6/8.

## Results

### Structure determination of the closed XPR1

To gain molecular insights into how KIDINS220 regulates XPR1, we first predicted the structure of XPR1 and KIDINS220 by AlphaFold2-multimer, which showed that the N-terminal ankyrin repeat domain (ARD, residues 1-432) of KIDINS220 interacts with residues 631-655 and the SPX domain of XPR1 (Supplementary Fig. 1a). To confirm this interaction, we further predicted the structures of XPR1 in complex with the ARD of KIDINS220, which showed a highest DockQ score of 0.81 with the same binding pattern as XPR1 and full length KIDINS220 (Supplementary Fig. 1b). To verify this, we co-expressed and purified the XPR1-KIDINS220 (1-432) complex, which showed XPR1 and KIDINS220 (1-432) form a stable complex (Supplementary Fig. 1c). Subsequently, we performed single-particle cryo-EM analysis and determined its structures at an overall resolution of 3.32 Å in the absence of 10 mM KH_2_PO_4_ and 3.36 Å in the presence of 10 mM KH_2_PO_4_ (Supplementary Fig. 2). Strikingly, both structures adopt the closed state conformations without bound phosphate in the binding pocket (hereafter referred to as XPR1^C^ for XPR1^C1^ and XPR1^C2^). The resulting model reveals only the transmembrane (TM) domain of XPR1, which assembles as a homodimer (Fig. 1a, b). The search in the Dali server for similar structures returned no significant hit. The TMD of XPR1 is composed of 10 helices (TM1-10) and the dimer interface is primarily formed by TM1 and TM3 from both protomers. Additionally, several lipid molecules contribute to the stabilization of this interface (Fig. 1b). The H2 helix of XPR1 is positioned between TM4 and TM5 and lies parallel to the membrane plane (Fig. 1b, c). The N-terminal SPX domain (residues 1 to 255), the C-terminal residues (residues 627 to 696) of XPR1 and KIDINS220 (1-432), are entirely absent in these structures. This may potentially be attributed to their significant dynamics.

**Fig. 1.**
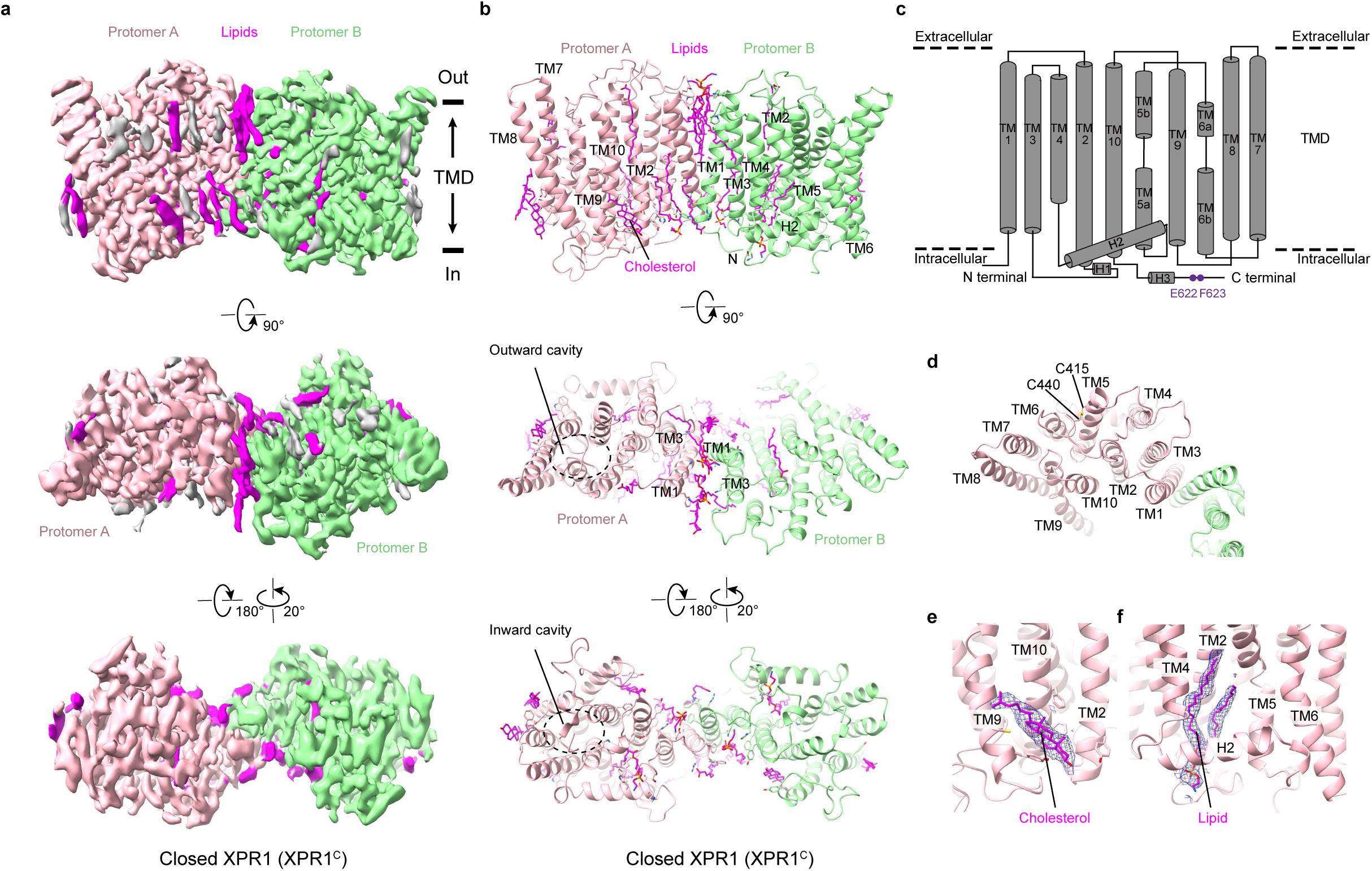
Structure of human XPR1 in the closed state. **a**, Cryo-EM maps of the closed state XPR1 (XPR1^C^) from side view, extracellular view and cytosolic view. The two protomers are colored in light pink and light green. Lipids are shown in magenta. The approximate boundaries of the phospholipid bilayer are indicated as thick black lines. **b**, Structural model of XPR1^C^ shown in cartoon representation. The two protomers and lipids are colored as in **a**. The outward cavity and intracellular cavity are labelled from extracellular view and cytosolic view, respectively. **c**, Schematic diagram of the secondary structure features of XPR1’s TMD. The Glu622/Phe623 motif near the C-terminal of XPR1 is labeled in purple. **d**, The potential disulfide bond between C415 and C440. **e**, Cryo-EM density map of the cholesterol at the pocket formed by TM2, TM9 and TM10, which is shown as blue meshes. **f**, Cryo-EM density map of the lipid at the pocket formed by TM2, TM4, H2 and TM5, which is shown as blue meshes.

In addition, Cys415 and Cys440 form a disulfide bond to stabilize TM6 (Fig. 1d). A cholesterol molecule is located at a groove formed by TM2 and TM9-10, which may stabilize the intracellular half of TM9 (Fig. 1b, e). We also observed another lipid resembles phosphatidyl ethanolamine with an aliphatic tail sticking into a pocket formed by TM2 and TM4-5, with the hydrophilic head locating underneath H2 (Fig. 1b, f). Overall, XPR1 forms a closed homodimer with abundant lipids contributing to its assembly.

### InsP6 facilitates the binding of SPX to transmembrane domain (TMD) in the presence of KIDINS220

Recent studies have reported that InsP8 binds to the N-terminus of XPR1, facilitating its phosphate export function^29,33^. This finding prompted us to investigate the mechanism by which InsP8 regulates this process. Due to the challenges associated with synthesizing InsP8, we employed InsP6 as a viable substitute. Subsequently, we elucidated the structures of the XPR1-KIDINS220 (1-432) complex in the presence of 10 mM InsP6 under two distinct conditions (Supplementary Fig. 3). The first structure was determined at an overall resolution of 3.68 Å in the presence of both 10 mM InsP6 and 10 mM KH_2_PO_4_. The second structure, resolved at 3.8 Å, was obtained in the presence of 10 mM InsP6 but in the absence of KH_2_PO_4_. Intriguingly, the two structures revealed distinct conformational states. The first structure (named as XPR1^OUT^) exhibited an outward-open conformation, with several well-defined electron densities resembling phosphate molecules (Fig. 2a, b). In contrast, the second structure (named as XPR1^C_SPX^) adopted a substrate-free closed state (Fig. 2c, d).

**Fig. 2.**
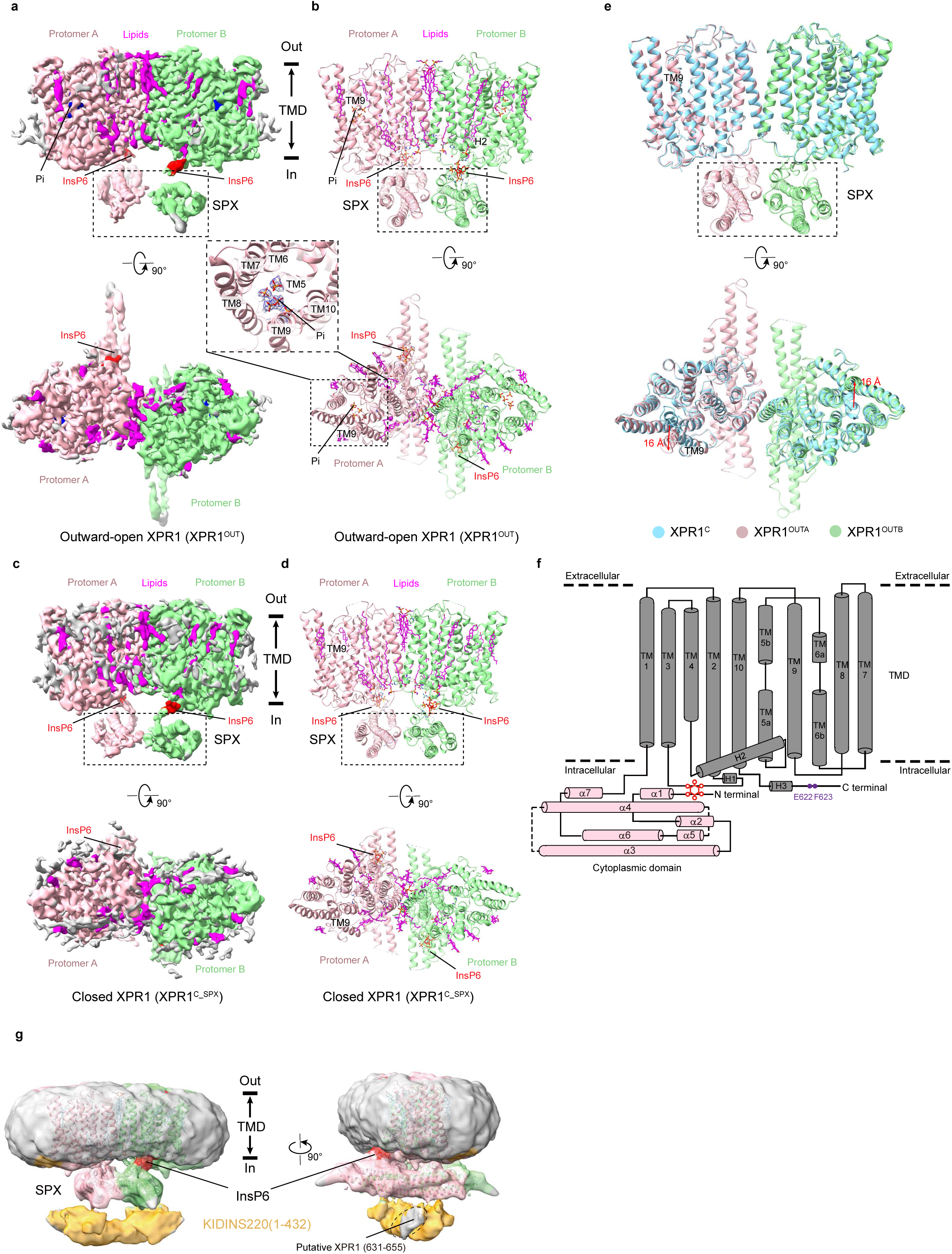
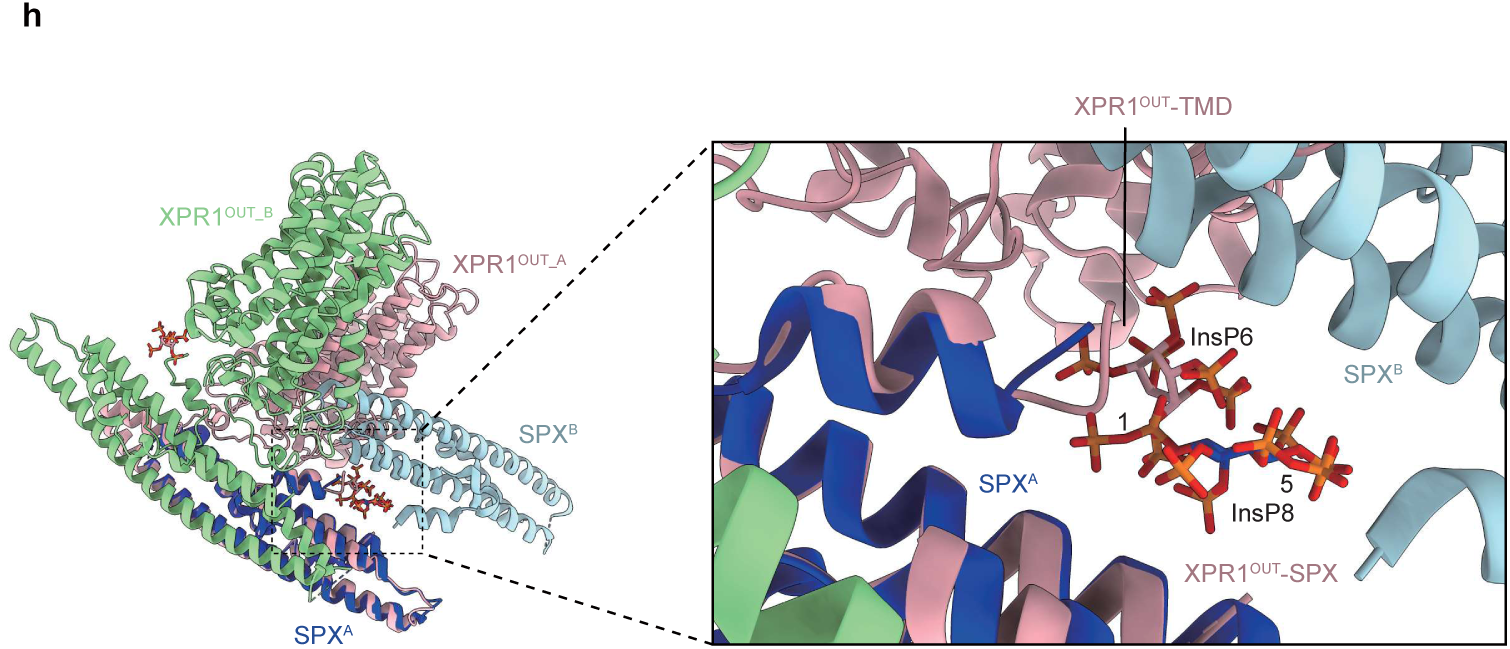
Structure of human XPR1 in the outward-open state. **a**, Cryo-EM maps of the outward-open state XPR1 (XPR1^OUT^) from side view and extracellular view. The two protomers are colored in light pink and light green. Lipids are shown in magenta. InsP6 molecules are shown in red. The approximate boundaries of the phospholipid bilayer are indicated as thick black lines. The densities of Pi are colored in blue. **b**, Structural model of XPR1^OUT^ shown in cartoon representation. The two protomers and lipids are colored as in **a**. The Pi and InsP6 molecules are labelled and shown in sticks. Cryo-EM density map of the Pi are shown as blue meshes as an insert. **c**, Cryo-EM maps of the closed state XPR1 with SPX (XPR1^C_SPX^) from side view and extracellular view. The two protomers are colored in light pink and light green. Lipids are shown in magenta. InsP6 molecules are shown in red. The approximate boundaries of the phospholipid bilayer are indicated as thick black lines. **d**, Structural model of XPR1^C_SPX^ shown in cartoon representation. The two protomers and lipids are colored as in **c**. The InsP6 molecules are labelled and shown in sticks. **e**, Structural comparison of XPR1^C^ (cyan) and XPR1^OUT^ (light pink and light green), superimposed based on the TMDs. **f**, Schematic diagram of the secondary structure features of full-length XPR1^OUT^. The TMD and the SPX domain are presented in gray and pink. The Glu622/Phe623 motif near the C-terminal of XPR1 is labeled in purple. InsP6 is labelled in red. **g,** Schematic diagram of the binding between KIDINS220 (1-432) and XPR1 in the outward-open state conformation. The XPR1^OUT^ atomic model was superimposed into the XPR1^OUT^ state density map after additional two rounds of 3D classification at low counter level. The two protomers of XPR1 and InsP6 molecules are presented in light pink, light green and red, respectively. The electron density maps of XPR1 and KIDINS220 at low counter level are shown in transparent mode and colored gray and orange, respectively. The binding pattern is based on the Alphafold2 prediction result presented in Supplementary Fig. 1a, b. **h**, Superimposition of our XPR1^OUT^ structure with the previously reported XPR1-SPX-1,5-InsP8 crystal structure by aligning one of the SPX domains. The InsP6 molecules and the protomer of XPR1^OUT_A^ were colored in light pink and the protomer of XPR1^OUT_B^ in light green. The InsP8 molecule and the SPX^A^ protomer of XPR1-SPX-1,5-InsP8 complex were colored in dark blue, and the SPX^B^ protomer in light blue.

Compared to the substrate-free closed state XPR1^C^, the most obvious differences are a central kink (∼45°) in TM9 and the densities for SPX domain of XPR1 and KIDINS220 (1-432) appear in the outward-open structure (Fig. 2e). The kink in TM9 created an extracellular entrance, extending from the membrane to the pore lumen within each protomer. When the scaffold domain of XPR1 was used as the reference to superimpose the TMD structure of XPR1^OUT^ onto that of XPR1^C^, the Cα atom of Thr582 in TM9 is observed to translate a distance of 16 Å (Fig. 2e). The SPX domain of each protomer is arranged parallel and in close proximity to the TMD, with the InsP6 binding site situated near the H2 helix (hereafter referred to as conformation 1 of the SPX domain) (Fig. 2b, 2d and 2f). KIDINS220 (1-432) binds on the opposite side of this arrangement and can only be discernible at low contour levels in the electron density map (Fig. 2g). Additionally, we superimposed our XPR1^OUT^ structure with the previously reported crystal structure of XPR1-SPX–1,5-InsP8 (PDB ID 8TYV)^33^. We found that the overall binding sites of InsP6 and InsP8 are similar, but the InsP6 molecules in our XPR1^OUT^ structure bind both with the SPX domain and the TMD and shift slightly toward the TMD (Fig. 2h). In summary, our structural analysis reveals that InsP6 simultaneously binds to both of the SPX domain and the TMD, thereby facilitating the interaction between these two domains.

### InsP6-binding site

Our structure shows that InsP6 binds to both the SPX domain and the peripheric juxtamembrane sides of XPR1, and the latter is formed by the intracellular loop between TM2 and TM3, TM4 and H2 (Fig. 2f, 3a and 3b). In particular, the positively charged Arg308 from the loop between TM2-TM3, Lys364 and Lys369 from the loop between TM4-H2 form potential salt bridge bonds with the negatively phosphate groups of InsP6 (Fig. 3c). Lys2, Lys158, Lys161, Lys162 and Lys165 from the SPX domain forms extensive salt bridge bonds with the negatively phosphate groups of InsP6 (Fig. 3c). Overall, the SPX domain and the TMD form a highly positive pocket and the negatively charged InsP6 functions as an intra-molecular glue to lock the orientation of the SPX domain and the TMD (Fig. 3b). Given that InsP8 carries more negative charges than InsP6, we hypothesized that InsP8 would more effectively facilitate the binding between the SPX domain and the TMD. We performed molecular dynamics (MD) simulations and verified this hypothesis. The results showed that Lys2, Lys158, Lys161, Lys162 and Lys165 from the SPX domain and Arg308, Lys364, Lys369 from the TMD clamp InsP8 tightly (Fig. 3d). These residues are all highly conserved (Supplementary Fig. 4). We further verified the function of this interaction by phosphate efflux assay. Compared to cells expressing wild-type (WT) XPR1, the efflux of radiolabeled ^32^P-phosphate was significantly reduced in cells overexpressing XPR1 mutants K2A, K158A, K162-165A, R308A, K364A, and K369A (Fig. 3e-g). Among these mutants, K2A, K158A, K162-165A and K369A showed most markedly reduced efflux of radiolabeled phosphate. Subsequent Pi efflux assays, which were conducted on the triple-mutation XPR1 mutants within the InsP6 binding pocket located in the SPX domain (XPR1-K158A-K162A-K165A) or the TMD (XPR1-R308A-K364A-K369A), revealed a more strikingly diminished Pi export activity (Fig. 3h, i). These results revealed that the InsP6/8 facilitated binding of the SPX domain and the TMD plays important roles in the phosphate export activity of XPR1.

**Fig. 3.**
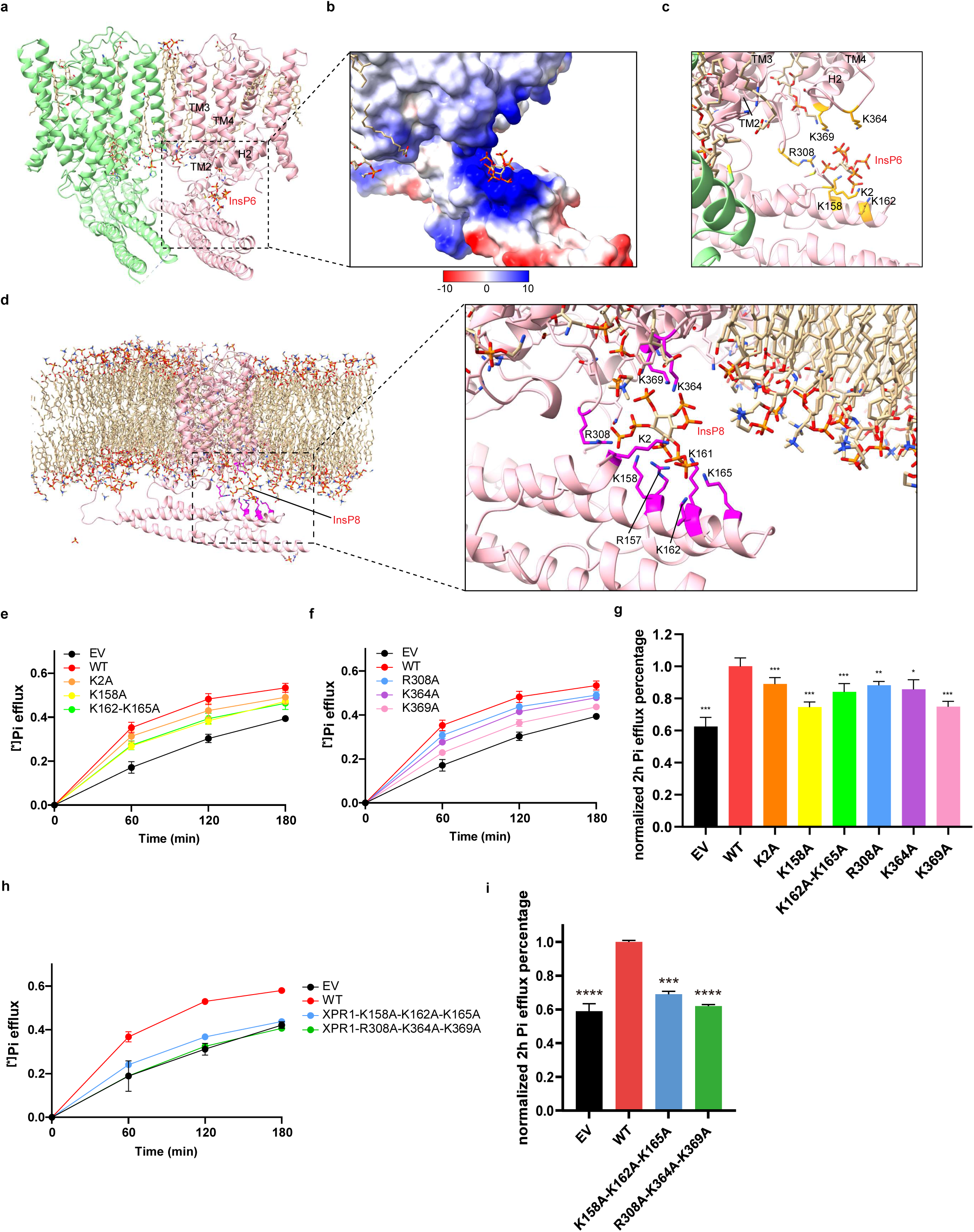
Inositol pyrophosphate bound to the SPX domain and peripheric juxtamembrane sides. **a**, One InsP6 molecule bound to a XPR1 protomer is presented in stick representation, and the lipid molecules are shown likewise, with carbon backbones colored in yellow. **b**, The electrostatic surface around InsP6. InsP6 molecule is displayed as sticks. **c**, Interactions between InsP6 and the surrounding residues from the SPX domain and the TMD. **d**, The binding configurations of InsP8 in three trajectories extracted from one of 1000 ns simulations. Interactions between InsP8 and the surrounding residues (magenta) from the SPX domain and TMD are shown in sticks. **e**,**f**, ^32^Pi efflux of HEK293T cells transfected with empty vector and constructs of WT-XPR1 or XPR1 mutants of residues from the InsP6 binding pocket in SPX domain (**e**) or TMD (**f**). Samples were taken and measured at indicated time points and the ratios of ^32^Pi against the total cellular ^32^Pi were calculated. Data are shown as mean ± s.d.; Data are shown as mean ± s.d.; n≥3 biologically independent experiments. **g**, ^32^P efflux percentages of EV and XPR1 mutants in (**e** and **f**) at 2 h normalized against the values of WT-XPR1 obtained under identical conditions from the same experimental batch. Data are shown as mean ± s.d.; n≥3 biologically independent experiments. *P* values (ns ≥ 0.05, *P < 0.05, *\*\*P <* 0.01, *\*\*\*P <* 1 × 10^-3^, *\*\*\*\*P <* 1 × 10^−4^) were obtained by unpaired *t* test with Welch’s correction between EV, XPR1 mutants and WT-XPR1. **h**, ^32^Pi efflux of HEK293T cells transfected with empty vector and constructs of WT-XPR1 or XPR1 triple-mutation mutants of residues from the InsP6 binding pocket in the SPX domain or the TMD. Samples were taken and measured at indicated time points and the ratios of ^32^Pi against the total cellular ^32^Pi were calculated. Data are shown as mean ± s.d.; n≥3 biologically independent experiments. **g, i**, ^32^P efflux percentages of EV and XPR1 mutants in **h** at 2 h normalized against the values of WT-XPR1 obtained under identical conditions from the same experimental batch. Data are shown as mean ± s.d.; n≥3 biologically independent experiments. *P* values (ns ≥ 0.05, *\*P* < 0.05, *\*\*P* < 0.01, *\*\*\*P* < 1 × 10 ^3^, *\*\*\*\*P* < 1 × 10 ^4^) were obtained by unpaired *t* test with Welch’s correction between EV, XPR1 mutants and WT-XPR1.

### A C-terminal plug-in loop blocks the intracellular cavity

We observed a C-terminal plug-in loop inserts into the inward-facing cavity of the XPR1 protomer. The position of the plug-in loop inside the cavity is stabilized by the interactions between the cytosolic extension of TM10 and the cytosolic cavity formed by TM5-8 and TM10 (Fig. 4a). Glu622 and Phe623 in the plug-in loop form critical interactions with the TMD. Based on these observations, we have designated this sequence as the “Glu622/Phe623 motif” (Fig. 4a). As illustrated in Fig. 4a, Glu622 forms a salt bridge with Arg472 from TM7. Phe623 engages in π-π stacking interactions with Phe391 and Phe394 from TM5, while also participating in a cation-π interaction with Arg466 from TM6. Notably, all these residues are highly conserved (Supplementary Fig. 4). To investigate the potential inhibitory role of the Glu622/Phe623 motif on XPR1’s phosphate export activity, we engineered a double mutant by substituting both residues with alanine (E622A/F623A). Remarkably, this double mutant exhibited a significantly enhanced rate of phosphate export activity compared to the WT-XPR1 (Fig. 4b, c). These findings provide compelling evidence for an auto-inhibitory function of the Glu622/Phe623 motif in regulating XPR1 activity.

**Fig. 4.**
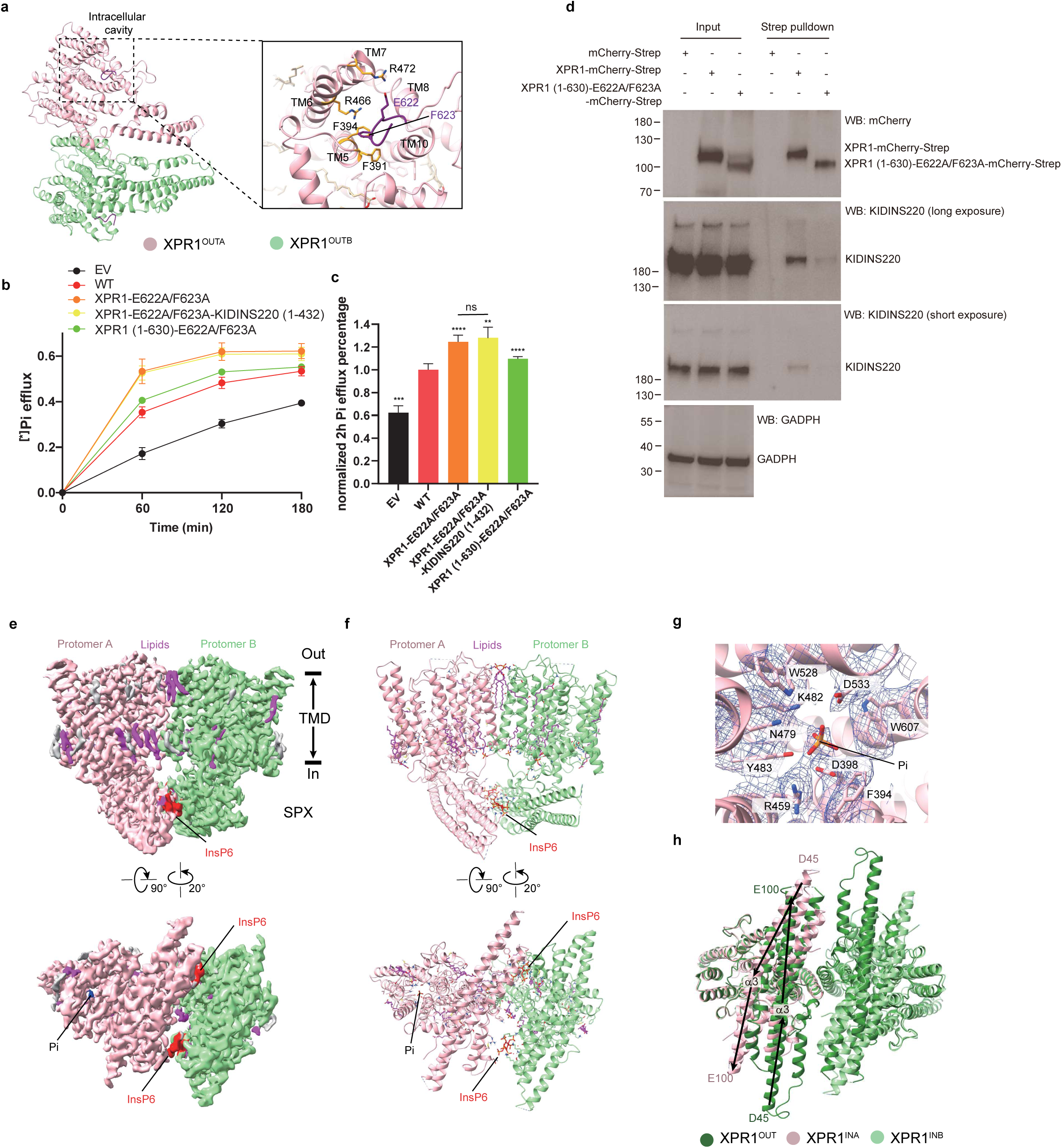
The C-terminal Glu622/Phe623 motif blocks the intracellular cavity. **a**, The Glu622/Phe623 motif and the cavity-forming helices from the TMD, i.e., TMs 5, 6, 7, 8, and 10. The Glu622/Phe623 motif is colored in purple, and shown as sticks. Interactions between the Glu622/Phe623 motif and the cavity-forming helices are shown. Putative interactive residues are displayed as sticks. **b**, ^32^Pi efflux of HEK293T cells transfected with empty vector and constructs of WT-XPR1 or XPR1 mutants of residues from the C-terminal plug-in loop. Data are shown as mean ± s.d.; n≥3 biologically independent experiments. **c**, ^32^P efflux percentages of EV and XPR1 mutants in (**b**) at 2 h normalized against the values of WT-XPR1 obtained under identical conditions from the same experimental batch. Data are shown as mean ± s.d.; n≥3 biologically independent experiments. *P* values (ns ≥ 0.05, \**P* < 0.05, **P < 0.01, ***P < 1 × 10^-3^, ****P < 1 × 10^-4^) were obtained by unpaired t test with Welch’s correction between EV, XPR1 mutants and WT-XPR1 or between XPR1-E622A/F623A mutant and XPR1-E622A/F623A-KIDINS220 (1-432) mutant. **d**, Western blot analysis of the binding capacity between KIDINS220 and either WT-XPR1 or XPR1 (1-630)-E622A/F623A. Input samples and Strep beads pulldown samples from HEK293T cells transfected with EV, WT-XPR1 and XPR1 (1-630)-E622A/F623A plasmids were subjected to SDS-PAGE and then probed with anti-GADPH, anti-mCherry and anti-KIDINS220 primary antibodies. Regarding the results of the KIDINS220 bands, both long and short exposure times were utilized to more effectively illustrate the binding capacity. **e**, Cryo-EM maps of the inward-facing XPR1 (XPR1^IN^) from side view and intracellular view. The two protomers are colored in light pink and light green. Lipids are shown in magenta. InsP6 molecules are shown in red. The approximate boundaries of the phospholipid bilayer are indicated as thick black lines. **f**, Structural model of XPR1^IN^ shown in cartoon representation. The two protomers and lipids are colored as in **e**. The Pi and InsP6 molecules are labelled and shown in sticks. **g**, Cryo-EM density map of the Pi and the surrounding residues at the intracellular cavity are shown as blue meshes. **h**, Structural comparison of XPR1^OUT^ (green) and XPR1^IN^ (light pink and light green), superimposed based on the TMD. The directions of the α3 helix backbones are indicated by arrows. The amino acid residues at the N- and C- ends of the α3 helices are labeled. Taking the α3 helix as an example, the SPX domain rotates nearly 180 degrees.

Structural prediction analyses revealed an α-helix (spanning residues 631 to 655) in the extension of the Glu622/Phe623 motif, which is predicted to interact with the N-terminal region of KIDINS220 (residues 1-432) (Supplementary Fig. 1a, b). This observation led us to hypothesize that KIDINS220 (1-432) might facilitate the dissociation of the Glu622/Phe623 motif from the intracellular cavity of XPR1. The hypothesis is corroborated by our experimental findings that the mutant XPR1 (1-630)-E622A/F623A, which lacks the ability to block the intracellular cavity and shows significantly reduced binding capacity with KIDINS220 (Fig. 4d), exhibited enhanced phosphate export activity compared to WT-XPR1 (Fig. 4b, c). Moreover, we also proved that KIDINS220 protein is expressed in the HEK293T cells that we used for the Pi efflux assays through western blot analysis (Fig. 4d). As a result, the endogenously expressed KIDINS220 could bind with the remaining XPR1, so that the co-expressed KIDINS220 (1-432) has no significant effect on the Pi efflux of XPR1 or XPR1− E622A/F623A (Supplementary Fig. 5a, b and Fig. 4b, c).

To investigate whether E622A/F623A mutation of XPR1 could induce an inward-facing conformation, we determined the structure of XPR1-E622A/F623A at an overall resolution of 3.03 Å in the presence of 10 mM InsP6 and 10 mM KH_2_PO_4_ (Supplementary Fig. 5c-j). Notably, in contrast to the outward-open structure of XPR1 complexed with KIDINS220 (XPR1^OUT^), XPR1-E622A/F623A adopts an inward-facing conformation with phosphate bound in the intracellular cavity (hereafter referred to as XPR1^In^) (Fig. 4e-g). Strikingly, the SPX domains of XPR1 rotate about 180 degrees (Fig. 4h), and two molecules of InsP6 are located between the two SPX domains (hereafter referred to as conformation 2 of the SPX domain) (Fig. 4e, f), further stabilizing the overall architecture of XPR1. The presence of KIDINS220 in XPR1^OUT^ would likely hinder the SPX domains from aligning as observed in the XPR1^In^ structure. In summary, these results show that the Glu622/Phe623 motif in the XPR1 protein plays an auto-inhibitory role in regulating the phosphate export activity, while the KIDINS220 protein may facilitate the dissociation of the Glu622/Phe623 motif to modulate the function of XPR1.

### The putative phosphate ion transport pore of XPR1

Based on our structures in the inward-facing and outward-open state conformations, we can infer the putative pathway for phosphate ion transport (Fig. 5a, b). The phosphate ion transport pore of XPR1 can be divided into three critical regions from the extracellular side to the cytosol (Fig. 5c). At the extracellular side, Y443, Q452, W514 and W573 form additional constrictions at the bottom of the pore like a gate (Fig. 5d). Four positively charged residues from TM7, TM9 and TM10 (K482, R570, R603, and R604) protrude into the pore, forming a positively charged ring within the membrane (Fig. 5e). They are associated with other conserved residues from TM4, TM8, and TM10 (D398, N401, S402, Y483, T525, D529, E600, W607) mediating an intricate network of interhelical interactions. At the intracellular entrance, a positively charged ring is formed by F394, R459, R466, R472, H476, W528, D533 and R611, facing the pore (Fig. 5f). Overall, the XPR1 pore is lined with highly conserved and predominantly positively charged residues (Supplementary Fig. 4).

**Fig. 5.**
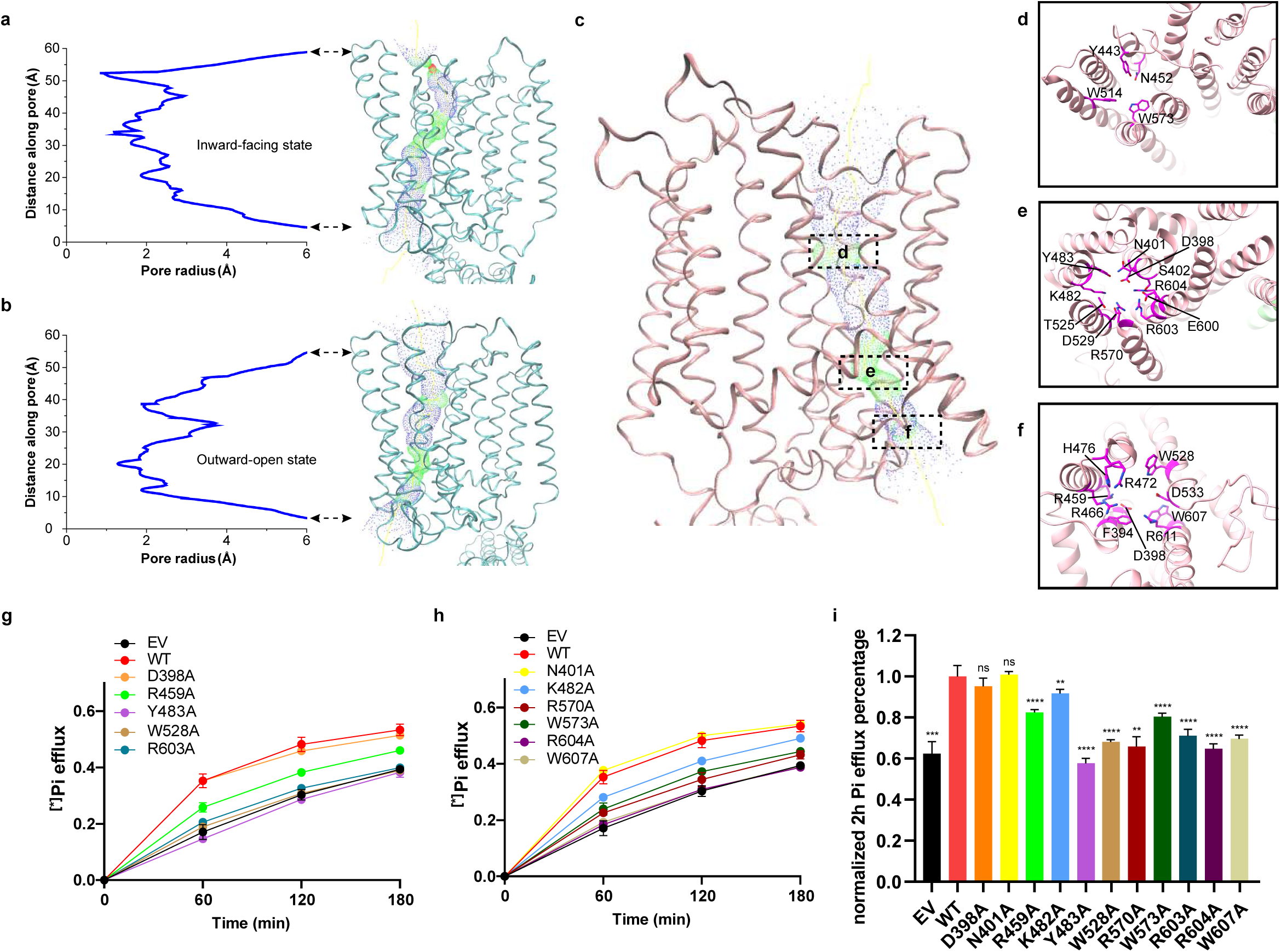
Putative phosphate ion pore. **a**, **b**, Plots of the pore radius as a function of the pore axis of XPR1^IN^ (**a**) and XPR1^OUT^ (**b**). The pore-lining surfaces were calculated using the program HOLE2^57^ and depicted on ribbon models of the XPR1^IN^ (**a**) and XPR1^OUT^ (**b**). **c**, The phosphate ion transport pore of XPR1^OUT^. The critical regions of the pore are labeled with dotted boxes. **d**, **e**, **f**, The top views (**d** and **e**) and the bottom view (**f**) of the cross sections through the ion-transporting pore at indicated positions in **c**. The ribbon is colored as in **c**. The conserved pore-lining residues are shown in sticks and colored in magenta. **g**,**h**, ^32^Pi efflux of HEK293T cells transfected with empty vector and constructs of WT-XPR1 or XPR1 mutants of residues from the phosphate ion transport pore. Data are shows as mean ± s.d.; n≥3 biologically independent experiments. **i**, ^32^P efflux percentages of EV and XPR1 mutants in (**g** and **h**) at 2 h normalized against the values of WT-XPR1 obtained under identical conditions from the same experimental batch. Data are shown as mean ± s.d.; n≥3 biologically independent experiments. *P* values (ns≥0.05, *P < 0.05, *\*\*P <* 0.01, *\*\*\*P <* 1 × 10^-3^, *\*\*\*\*P <* 1 × 10^−4^) were obtained by unpaired *t* test with Welch’s correction between EV, XPR1 mutants and WT-XPR1.

Together, the structural feature of XPR1 likely contributes to its function as an anion transporter. The findings above suggest that the transported anions interact with the pore-lining charged residues, thus participating in transporter regulation. To test the residues responsible for phosphate export, we mutated residues D398, N401, R459, K482, Y483, W528, R570, W573, R603, R604 and W607 to alanine. Among these mutants, Y483A and W528A showed the most dramatic reductions in phosphate export activity (Fig. 5g-i). Our structural analysis revealed that both Y483 and W528 are located within the pore and interact with residues forming the positively charged ring, which can maintain the stability of the pore (Fig. 5e, f). Disruption of these key interactions likely leads to the collapse of the pore, thereby impairing the phosphate export function of XPR1.

The structural features of the XPR1 pore, including the positively charged rings and gate-like constrictions, suggest a putative pathway for phosphate ion transport, which is supported by the functional impact of mutations in key pore-lining residues.

### The proposed model for XPR1 regulation

Based on our structural data and functional studies, we put forward a working model elucidating the manner in which KIDINS220 and inositol pyrophosphate modulate the Pi export activity of XPR1 (Fig. 6). In our proposed regulatory model, elevated cytosolic phosphate concentration triggers an increase in intracellular InsP8 levels, which subsequently promotes InsP8’s binding to the SPX domains of XPR1. In the absence of KIDINS220, the SPX domains of XPR1 adopt an antiparallel arrangement in conformation 2. In this configuration, InsP8 molecules bind at the interface between the two SPX domains, further stabilizing this conformation. This stabilized state appears to impede the efficient release of the C-terminal plug-in loop, consequently maintaining XPR1 in an inactive state. However, in the presence of KIDINS220, the SPX domain shifts to another antiparallel arrangement in conformation 1, which is rotated about 180 degrees relative to conformation 2. In this state, the binding of InsP8 between the SPX domain and the TMD lowers the barrier of the disassociation of the C-terminal plug-in loop from the intracellular cavity, thus initiating the phosphate export activity of XPR1. Therefore, both InsP8 and KIDINS220 play crucial roles in modulating XPR1’s phosphate efflux activity.

**Fig. 6.**
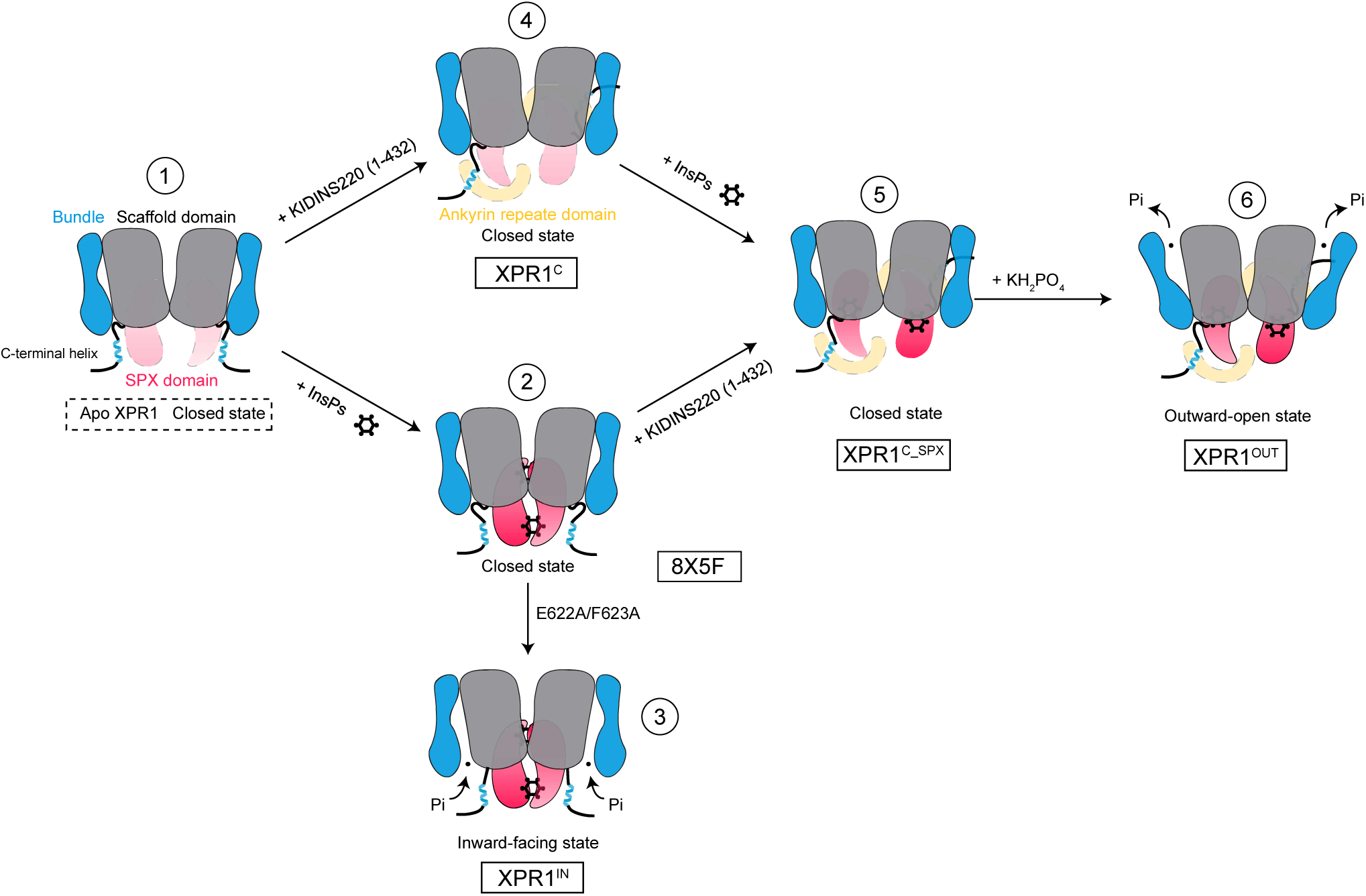
Putative activation model for the InsPs and KIDINS220 modulated phosphate exporter in XPR1. A schematic representation of the proposed XPR1 activation mechanism. Under non-activating conditions, the TMD is locked in a closed state (**state 1**). When KIDINS220 is absent, elevated InsPs in the cytoplasm bind between the two SPX domains of XPR1, locking XPR1 in a closed conformation (**state 2**). Mutation of the Glu622/Phe623 motif which blocks the intracellular cavity induces XPR1 to transition to an inward-facing conformation (**state 3**). In the first three states, the flexible C-terminal helix (631-655) resides near the SPX domain. In the presence of KIDINS220, the C-terminal helix binds with KIDINS220 (1-432) and induces further conformational change of the SPX domain possibly due to steric hindrance. Although the dynamics of the SPX domain are too strong to observe its specific orientation, subsequent results indicate that its conformation undergoes a 180-degree flip compared to **state 2**, with XPR1 still remaining in a closed conformation (**state 4**). InsPs bind at the interface between the SPX domain and the TMD, maintaining XPR1 in a closed conformation. However, this binding enhances the dynamics of the Glu622/Phe623 motif, thereby eliminating one of the barriers to activation (**state 5**). Upon the introduction of substrate phosphate ions, they promptly bind to XPR1, thereby inducing the conformational alteration of XPR1 into an outward-open state. This conformational change, in turn, promotes the export of these phosphate ions (**state 6**). Although the two halves of the XPR1 dimer in different states were depicted with the same behavior in our figure drawing, we believe that their actual functional states might not be in synchronization. **States 3 to 6** were observed in this study. **State 2** is derived from PDB ID: 8X5F, and **state 1** is hypothesized.

## Discussion

In these years, several studies have investigated the impacts of the SPX domains and inositol pyrophosphate on the phosphate export activity of XPR1. Wild et al.^27^ determined the crystal structures of the SPX domains of diverse eukaryotic proteins and verified the significant roles of many conserved positively charged amino acids in the SPX domains. Li et al.^29^ firstly identified 1,5-InsP8 as the relevant regulation ligand of XPR1 in its phosphate efflux activity *in vivo*. Moreover, recently Li et al.^33^ has reported the crystal structure of XPR1-SPX-1,5-InsP8 and established that XPR1 may function as a hetero-dimer with PiT1 *in vivo*. Nevertheless, the specific molecular evidence regarding how XPR1 achieves its phosphate export activity and how InsP8 facilitates this process continues to be indistinct. In this study, we have investigated the phosphate transport properties of XPR1 using cell-based phosphate transport assays and determined cryo-EM structures of the human XPR1-KIDINS220 complex in the substrate-free closed states, and the phosphate-bound outward-open state. Our work provides insights into the architecture, phosphate binding and conformational landscapes of the phosphate exporter XPR1-KIDINS220 complex. Through multiple lines of evidence, we conclude that both inositol pyrophosphate and KIDINS220 are essential for the substrate transport activity mediated by XPR1.

We characterized the functional importance of several aromatic and positively charged residues within the substrate-binding site that participate in ligand coordination and transport. Based on our structural findings and MD simulation data, we propose a “rocking-bundle”, alternating-access mechanism to describe the transport cycle of phosphate export mediated by XPR1. In this proposed transport cycle model, the inward-facing conformation of XPR1 represents the state in which phosphate ions bind from the intracellular space. Phosphate-induced conformational changes then drive the transporter towards its outward-open state, allowing phosphate ions to be released into the extracellular space. After the release of phosphate, XPR1 will experience further conformational alterations to assume the inward-facing state conformation, thus restarting the transport cycle. Considering that phosphate ions are transported from the intracellular compartment to the extracellular space, and given that the concentration of potassium ions is greater within the cell compared to the exterior, we postulate that the driving force for phosphate ion transport might be the potassium ion concentration gradient, which requires further investigation.

Regarding other structural biology studies on XPR1, Jiang et al.^34^ reported the cryo-EM structures of XPR1^Open^ and XPR1^Closed^, which were obtained through 3D classification of apo XPR1, along with the structure of XPR1^InsP6^. Additionally, they carried out malachite green phosphate assays under Pi starvation conditions to investigate the roles of key residues on the Pi export activity of XPR1. Through structural analysis and functional results, they suggested a channel-like working model in which InsPs (inositol polyphosphates) serves to stabilize the SPX domains. The C-loop turns to bind the SPX domains, which consequently leads to the opening of the intracellular Pi gate. However, in their structure of XPR1^InsP6^, XPR1 was not observed in an open state, but rather in a closed state. A careful examination of the electron density in their XPR1^InsP6^ structure (PDB ID: 8X5F and EMDB: EMD-38068) reveals that the C-loop not only has a conformation bound to the SPX domain, but also shows clear density binding to the intracellular cavity, which deviates from their proposed working model (Supplementary Fig. 6a, b).

Cao et al.^35^ reported the cryo-EM structure of XPR1-IP_6_/PPF, which is analogous to the XPR1^InsP6^ structure reported by Jiang et al.^34^, and the structure of XPR1-IP_7_, where InsP7 binds between the SPX domain and the TMD of one XPR1 protomer. Additionally, they captured intermediate conformations of XPR1 in the XPR1-IP_6_/WO_4_ structure. Through structural studies and electrophysiological analyses, they proposed a model in which XPR1 acts as an inositol pyrophosphate activated channel. In contrast, our study employed the radioisotope ^32^P efflux assay to investigate the Pi export activity over longer timescales under physiological conditions. Moreover, KIDINS220 was introduced into our structural analysis, which has been previously reported to participate in the membrane-localization and Pi export activity of XPR1^31^. As a result, in comparison with the structures reported by Cao et al. and Jiang et al., the two XPR1 protomers in our XPR1^OUT^ structure (supplemented with KIDINS220 (1-432), 10 mM InsP6 and 10 mM KH_2_PO_4_) both adopt the outward-open state conformations. Moreover, the electron density of the phosphate ions was more distinct in our structures than in theirs. Although the TMD side of the InsP6 binding pocket in our XPR1^OUT^ structure is identical with that of the InsP7’s TMD-SPX site described by Cao et al., the binding pockets of InsP6/7 in the SPX side and the orientation of the SPX domains vary from each other. Therefore, we believe that both KIDINS220 and inositol pyrophosphate contribute to the activation of XPR1.

KIDINS220 has been reported to be located at late endosomes and the plasma membrane^36^. And our results verified that KIDINS220 is essential for the phosphate export activity of XPR1. We propose that the phosphate export activity of XPR1 can only be properly manifested when the transporter is localized to the membrane of organelles that contain the KIDINS220 protein. This likely serves as a mechanism to prevent the excessive accumulation of phosphate within other cellular compartments. In the case of late endosomes and the plasma membrane, the presence of KIDINS220 allows the cell to utilize XPR1 to export phosphate into late endosomes or the extracellular space. This helps maintain appropriate phosphate levels in the cytosol and prevent its excessive buildup. It is remarkable that the peak values in our Pi efflux assays did not exceed 65%. The possible explanation could be the continuous sequestration of intracellular Pi into endosomes and its subsequent conversion to ATP through G3PDH and oxidative phosphorylation. This also indicates the potential role of endosomes and KIDINS220 in cellular phosphate regulation. The synergistic regulation of XPR1 by both InsP8 and KIDINS220 appears to be a crucial control mechanism to ensure tight regulation of cellular phosphate homeostasis. By directing the localization and activity of the XPR1 phosphate exporter, the cell can effectively manage phosphate levels across different organelles and the extracellular environment. This intimate association of XPR1 function with particular organelle membranes, which is facilitated by KIDINS220, constitutes a significant regulatory mechanism in the preservation of the phosphate equilibrium within the cell.

In summary, our studies delve into the role and mechanism of the XPR1-KIDINS220 complex in cellular phosphate export. Through cell transport experiments and cryo-EM structural analysis, we revealed the structures, phosphate binding sites, and conformational changes of the XPR1-KIDINS220 complex. The study found that inositol pyrophosphates (particularly InsP8) and KIDINS220 are crucial for substrate transport by XPR1. A “rocking-bundle” alternating access mechanism was proposed to explain the phosphate export process mediated by XPR1. Moreover, multiple missense mutations in *XPR1* gene have been identified in patients diagnosed with PFBC^19,20,37–42^ or papillary thyroid carcinoma^43^. Through structural analysis, we have discovered that many of these mutations are strategically positioned along the phosphate ion transport pore or in close proximity to the Glu622/Phe623 motif and InsP8 binding site within the SPX domain (Supplementary Fig. 6). The distribution characteristic of the mutations is consistent with our proposed phosphate export mechanism of XPR1 and two of the residues (R459 and R570) have been investigated in our study. In brief, our findings provide new insights into the regulation of cellular phosphate homeostasis, which may have significant implications for the study and treatment of related diseases.

## Methods

### Constructs

Full-length human *XPR1* (Uniprot ID: Q9UBH6) gene and human *KIDINS220* (Uniprot ID: Q9ULH0) aa 1-432 fragment gene were cloned from human HEK293 cDNA and human brain tissue-derived cDNA, respectively. The gene fragments were then subcloned into the two ORFs of a modified pFastbac-dual-CMV-mCherry-6His-Strep-EF1α-eGFP (Empty vector, EV) plasmid. The full-length *XPR1* gene was also individually cloned into the CMV ORF of the EV plasmid. A series of point mutations of *XPR1* gene were introduced into the two plasmids mentioned above through KOD-plus-neo enzyme (TOYOBO) and Dpn1. The wild-type and mutant plasmids were subsequently used for protein expression and functional assays.

### Protein expression and purification

Bac-to-Bac baculovirus system (Invitrogen) and HEK293F (Gibco) cells were used for protein expression. Recombinant plasmids of wild-type XPR1-KIDINS220 (1-432), wild-type XPR1 and XPR1-E622A/F623A mutant were transfected into DH10Bac strains to produce bacmids, respectively. The extracted bacmids were transfected into SF9 cells using PEI to obtain the P0 generation baculoviruses. After two rounds of amplification, the P2 generation baculoviruses were used to infect the suspension-cultured HEK293F cells at the v/v ratio of 1:10. The cells were incubated at 37 °C in suspension supplemented with 1% (v/v) FBS and 5% CO_2_ for 12 h before the addition of 10 mM sodium butyrate and cultured for another 36 or 48 h before harvesting. The cells were then collected by centrifugation at 4000g for 10 minutes. The pellets were resuspended with 1*TBS buffer (140 mM NaCl, 3 mM KCl, 10% glycerol, 30 mM Tris-HCl, pH 7.5), then frozen in the liquid nitrogen and stored in −80 °C.

To separate the cell membranes, the thawed cells were supplemented with 1 mM PMSF and ultrasonically extracted for 2.5 min. The supernatants were collected after the centrifugation at 8000 g for 20 min and then ultracentrifuged at 100,000 g for 1 h to isolate the cell membranes. The cell membranes were ground thoroughly and then solubilized with dissolving buffer (150 mM NaCl, 20 mM Tris-HCl pH 7.4, 1% (w/v) n-dodecyl β-Dmaltoside (DDM, Anatrace), and 0.1% (w/v) cholesteryl hemisuccinate (CHS, Anatrace)), stirred at 100 rpm at 4 °C for 2 h. Insoluble debris was removed by centrifugation at 100,000 g for 30 min. The supernatants slowly flew through Ni-NTA beads at 4 °C after the addition of 20 mM imidazole. Nonspecifically bound protein was removed using 10 columns of elution buffer (150 mM NaCl, 50 mM HEPES, pH 7.4, and 0.01% (w/v) glyco-diosgenin (GDN, Anatrace)) in addition of 20 mM or 40 mM imidazole. Target proteins were subsequently washed off by 6 columns of elution buffer with 250 mM imidazole. After diluting the imidazole concentration, the proteins were incubated with Strep beads and then cut by TEV protease at 4 °C overnight to remove the mCherry-His6-Strep II tags. 3 columns of elution buffer were used to completely wash off the cleaved protein. For the InsP6 bound samples, 10 mM InsP6 was added at this step. Proteins were then concentrated and further purified by gel filtration (Superose 6 Increase 10/300 GL, GE Healthcare, USA). Peak fractions were collected and concentrated to nearly 10mg/mL for subsequent Cryo-EM sample preparation.

### Cryo-EM sample preparation and data acquisition

For the phosphate-bound samples, 10 mM Potassium dihydrogen phosphate (KH_2_PO_4_, pH 7.5) was added to the samples. 3 ul protein samples were loaded onto the glow-discharged Holey Carbon films (Au R1.2/1.3 300 mesh grids, quantifoil) or ANTcryo™ holy support films (M01-Au300-R1.2/1.3, Nanodim). The grids were automatically blotted with filter paper for 3 s under 100% humidity at 6 °C before being plunged into liquid ethane using Vitrobot Mark IV (Thermo Fisher Scientific).

Cryo-EM data were collected using 300 kV FEI Titan Krios electron microscopy (Thermo Fisher Scientific) equipped with K2 summit direct electron detector (Gatan) or FEI Falcon4 direct electron detector (Thermo Fisher Scientific). A magnification of 165kx with a calibrated pixel size of 0.821 Å (K2) or 0.808 Å (Falcon4) was used for movie acquisition. The dose rate and total dose of data collection was set to ∼ 8 e^-^ /pixel/s and ∼50 e^-^/Å^2^. A total exposure time of ∼5 s was used to film the movies which were dose-fractionated into 40 or 32 frames. SerialEM or Smart EPU Software were used to automatically acquire the data. For each protein sample condition, more than 3000 movie stacks were collected for further structural analysis.

### Cryo-EM data processing

The collected movie stacks were converted to micrographs through motion correction by MotionCor2^44^. The cryoSPARC software^45^ was used for subsequent data processing procedures. The corrected micrographs were imported and performed with Patch CTF estimation for CTF measuring and further image selection. The micrographs with inappropriate ice thickness or power spectra were excluded in the Manually Curate Exposures procedure. Blob picker and Inspect picks procedures were used to pick protein particles preliminarily. The particles were then extracted and conducted with 2D classification to generate good templates for further particle picking. Template picking and Topaz picking^46^ were used to improve and generate the final particle sets. After Remove Duplicates, the reserved particles went through 3-4 rounds of 2D classification to produce the cleanest “seed particles” for following particle retrieve. For the XPR1-E622A/F623A mutant in the presence of InsP6, two rounds of Ab-initio reconstruction were used here to get the seed particles. The particles selected in the first round of 2D classification were split into several particle sets based on the size of the seed particles. Different particle splits together with the seed particles were performed with parallel 2D classifications. The sum of the seed particles and the selected particles after removing duplicates were performed with 2 rounds of 2D classification to acquire the final particles for Ab-initio reconstruction. Typically, two or three initial models were generated. The volume and particles of each initial model were used to conduct Non-uniform refinement. To determine the density of the flexible SPX domains of XPR1 in the presence of KIDINS220 (1-432) and InsP6, the particles in the last procedure were divided into three classes in 3D classification and performed with Non-uniform refinements, respectively. Local resolution estimation procedures were employed to determine the resolution of each map.

### Model building, refinement and validation

The initial template of XPR1 was generated using Alphafold2^47^. The Cα backbone and side chains were carefully inspected and adjusted manually to conform with the cryo-EM map, utilizing Coot^48^ for visualization and manipulation. Multiple lipids were added to the model according to their corresponding densities in the maps. Depending on the different sample preparation conditions, InsP6 and putative water molecules, phosphate ions and potassium ions were incorporated into the models based on density similarity. The C-terminal region of 626-696 residues of XPR1 is not clearly visualized in all of the density maps due to its flexibility. Likewise, the flexible N-terminal 1-225 residues of SPX domains is also invisible in the conformations in which SPX domains are not stabilized by InsP6. Consequently, atomic models were not built for these highly flexible regions. The Real Space Refinement program of PHENIX^49^ was utilized to refine the atomic models against the cryo-EM maps. The final model statistics were validated and supplied by MolProbity^50^ in PHENIX, and are detailed in Supplementary Table 1. The sample conditions of the cryo-EM structures in our study are listed in Supplementary Table 2. Structural figures were generated using UCSF ChimeraX^51^.

### [^32^P] KH_2_PO_4_ efflux assays

HEK293T cells were seeded into six-well plates at an appropriate concentration. 2 μg of EV plasmid (expressing mCherry-Strep) or wild-type and mutant XPR1-mCherry-Strep plasmids were transfected using PEI and Opti-MEM into the HEK293T cells when density reached ∼50%. After 12 h, the medium was changed with 2 mL fresh DMEM supplemented with 10% FBS. The cells were further cultured for 24 h at 37 °C before phosphate efflux assay. The transfection efficiency of plasmids was first confirmed using a fluorescence microscope. The cells were incubated with 1 mL DMEM supplemented with 0.5 μCi/mL [^32^P] KH_2_PO_4_ for 20 minutes. The cells were then washed three times with 1 mL Pi-free DMEM and incubated with 1 mL phenol red-free DMEM supplemented with 1 mM KH_2_PO_4_ (pH 7.4). 100 μL of medium was taken at 1 h, 2 h and 3h. The rest of the medium was then removed and the cells were digested by trypsin. After centrifugation at 1000 g for 3 minutes, the cells were lysed for 30 minutes with PBS solution containing 1% Triton-X-100. Both the cell lysate and the previously collected aliquots of culture medium were mixed with 3 mL of liquid scintillation cocktail, and the ^32^P signal intensities were measured using a liquid scintillation counter. Pi efflux is described as the ratio of the total signal intensity in the culture medium at different time points relative to the total signal intensity including the cell lysate.

### Western blot analysis

In order to explore the interaction mode between KIDINS220 and XPR1, HEK293T cells were seeded into 10 cm dishes and transfected with 10 μg of EV, XPR1-mCherry-Strep, XPR1 (1-630)-E622A/F623A-mCherry-Strep plasmids using PEI and Opti-MEM, respectively. Following the replacement of the medium at 6 hours and an additional 30 hours of culturing, the cells were centrifuged and washed once with 1x PBS buffer, and then lysed with the lysis buffer (150 mM NaCl, 50 mM HEPES at pH 8.0, 1% NP-40, 1 mM PMSF and 1 mM protease inhibitor cocktail (Roche)) on ice for 30 min. The cell extracts were clarified by centrifugation at 13,000 g for 20 min at 4°C. The protein contents were quantified by bicinchoninic acid (BCA) analysis. 40 μg of proteins were taken as input samples and mixed with 5x loading buffer. Equal amounts of the residual proteins were incubated with 20 μL of Strep beads at 4 °C for 2 hours under slow rotation. The beads were washed three times with 500 μL of wash buffer (150 mM NaCl, 50 mM HEPES at pH 8.0, and 0.5% NP-40). Subsequently, the proteins were eluted with 30 αL of elution buffer (150 mM NaCl, 50 mM HEPES at pH 8.0, 50 mM biotin, and 1 mM EDTA) at room temperature for 30 minutes under slow rotation. The supernatant was separated and combined with 5x loading buffer to prepare co-IP samples. The input samples and co-IP samples (without boiling) were separated on 4– 20% gradient polyacrylamide gels and then transferred to PVDF membranes under constant current conditions. The PVDF membranes were cut according to molecular weight and then incubated overnight at 4 °C with the indicated primary antibodies, respectively. The following primary antibodies were utilized in this study: Anti-mCherry Tag Mouse Monoclonal Antibody (9D3) (Abbkine, ABT2080, diluted 1:5000), KIDINS220 Monoclonal antibody (Proteintech, 66748-1-Ig, 1:6000) and GAPDH antibody (Ray Antibody, RM2002, 1:3000). Subsequently, the membranes were washed four times in 1x TBST buffer (25 mM Tris at pH 7.5, 140 mM NaCl, 3 mM KCl and 0.1% Tween-20) and further incubated with Goat anti-Mouse IgG secondary antibody (Invitrogen, 31431, 1:5000) at room temperature for 45 minutes. The membranes were washed four times in 1x TBST buffer and then incubated with Western Blotting Luminol Reagent (Santa Cruz, sc-2048) to detect immunoreactive bands.

### All-atom molecular dynamics simulations

The outward-open state structure of XPR1-KIDINS220 (1-432) in complex with InsP6 was used as the initial structure for molecular dynamics simulations. The XPR1 was embedded into a lipid bilayer that simulated the components of the cell membrane (with 34% Chol, 23% POPC, 17% SM, 11% POPE and 8% POPS)^52^ using the CHARMM-GUI software package^53^. The system was subsequently solvated utilizing the TIP3 water model, and 150 mM Na^+^ and Cl ions were introduced to achieve overall charge neutrality. The CHARMM36m force field was utilized to model the behavior of proteins and lipids in the system^54^. To investigate the transport mode of phosphate, we introduced different quantities of phosphate ions into the phosphate transport channels of both monomers based on the density and residue electrostatic properties. Additionally, we reintroduced InsP6 or InsP8 near the corresponding density in the model to simulate different binding modes of inositol pyrophosphate molecules with the pocket. Unrestrained production simulations were performed with 1 μs time integration steps at a constant temperature of 310 K. Three independent simulation trajectories were conducted using the GROMACS software package^55^. Analyses were conducted using the GROMACS and the visual molecular dynamics (VMD) program^56^.

### Reporting summary

Further information on research design is available in the Nature Research Reporting Summary linked to this article.

## Data availability

The atomic model coordinates and cryo-EM maps have been deposited in the Electron Microscopy Data Bank (EMDB) and Protein Data Bank (PDB), respectively, under the accession codes: 9INE and EMD-60704 (XPR1^C1^), 9INF and EMD-60705 (XPR1^C2^), 9INH and EMD-60707 (XPR1^OUT^), 9IUC and EMD-60897 (XPR1^C_SPX^), 9ITG and EMD-60861 (XPR1^In^). Any additional information required to reanalyze the data reported in this paper is available from the Lead Contact upon request. Source data are provided with this paper.

## Acknowledgements

Cryo-EM data collection were supported by the Cryo-EM Platform of Peking University with the assistance of Dr. Xuemei Li, the Cryo-EM platform of Peking University Health Science Center with the assistance of Dr. Dandan Chen and Dr. Lihong Chen, and the cryo-EM facility from Shuimu BioSciences. Peng Zuo received training in cryo-EM sample preparation and data collection from Prof. Youdong Mao of the School of Physics, Peking University. We thank Dr. Hongjie Zhang and Prof. Junjie Hu from the Institute of Biophysics, Chinese Academy of Sciences for their assistance in the ^32^P efflux assays. This study is supported by grants to Y.Y. including the National Key Research and Development Program of China (2021YFA1300601, Y.Y.), National Natural Science Foundation of China (key grant 82030081 and 81874235, Y. Y.); Shenzhen High-level Hospital Construction Fund and Shenzhen Basic Research Key Project (JCYJ20220818102811024, Y.Y.). L.L. is supported by the National Natural Science Foundation of China (grants 32171224, L.L.). Y.Y. is the Scholar of the Lam Chung Nin Foundation for Systems Biomedicine. The MD simulations were supported by the high-performance computing platform of Peking University.

## Author contributions

Y.Y. and L.L. conceived the project. P.Z. established the constructs, purified proteins, performed biochemical experiments, and cell assays. P.Z. prepared cryo-EM grids and participated in the cryo-EM data collection. P.Z., and L.L. processed the cryo-EM data, built models and performed modelling and refinement. W.W., Z. D., G. Wang and Y. Y. helped with the data processing procedures. W.W. assisted in the Western Blot analysis. S.Y. assisted in the cryo-EM data collection. J.Z. participated in part of the protein purification. P.Z. and L.L. performed and analyzed the molecular dynamics simulation. P.Z., L.L. and Y.Y. wrote the manuscript with input from all authors.

---

**Supplementary Fig. 1.**
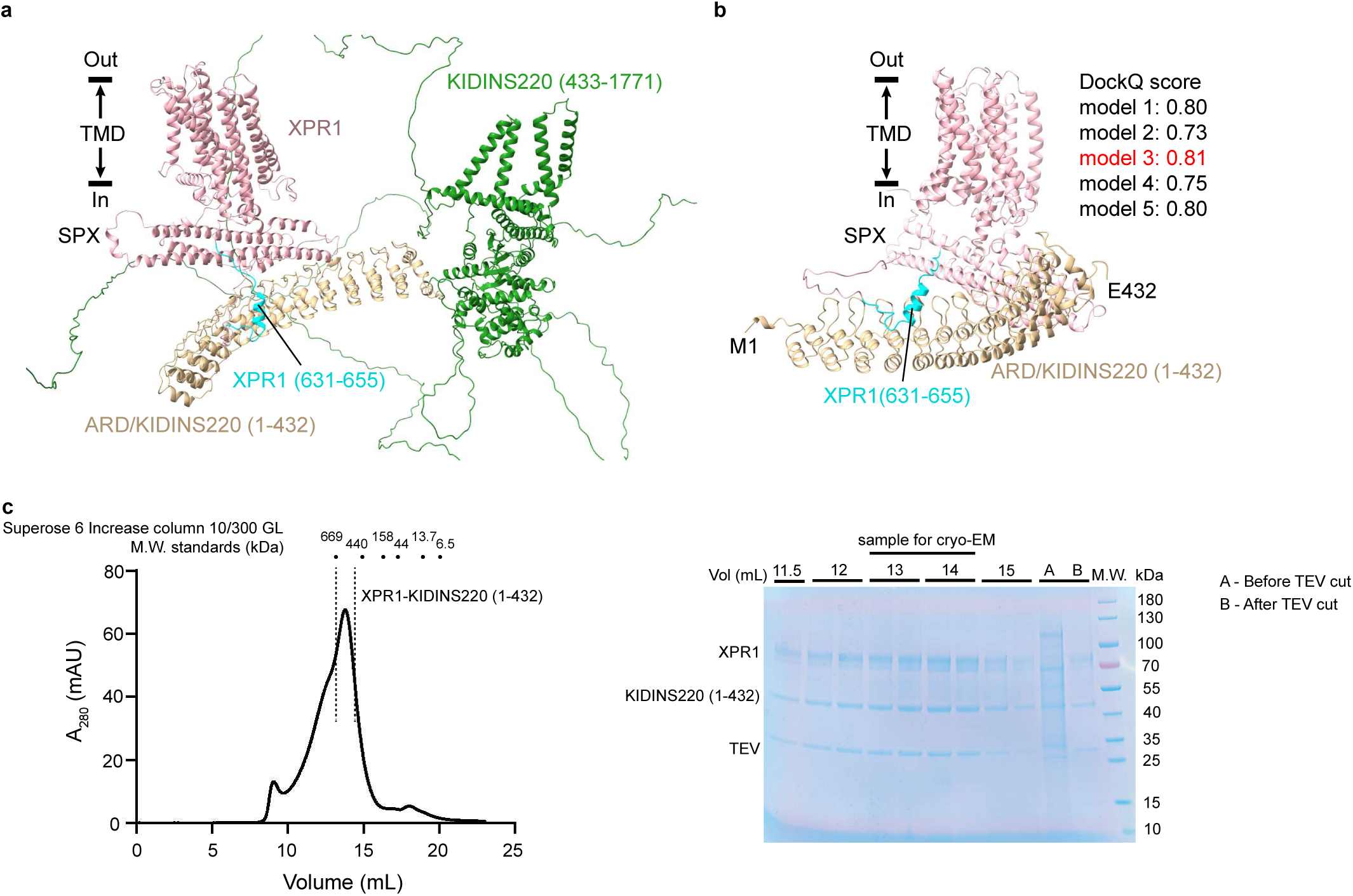
XPR1 stably interacts with KIDINS220 (1-432) **a**, AlphaFold2-multimer prediction of XPR1 and full-length KIDINS220. **b**, AlphaFold2-multimer prediction of XPR1 and KIDINS220 (1-432). The DockQ scores of the predicted models were shown in the figure. **c**, The size-exclusion chromatography (SEC) profile and SDS-PAGE gel of purified XPR1-KIDINS220 (1-432). Fraction within the dashed lines were concentrated for cryo-EM sample preparation. The molecular weight standards of the Superose 6 Increase column 10/300 GL were labeled on the SEC profile.

**Supplementary Fig. 2.**
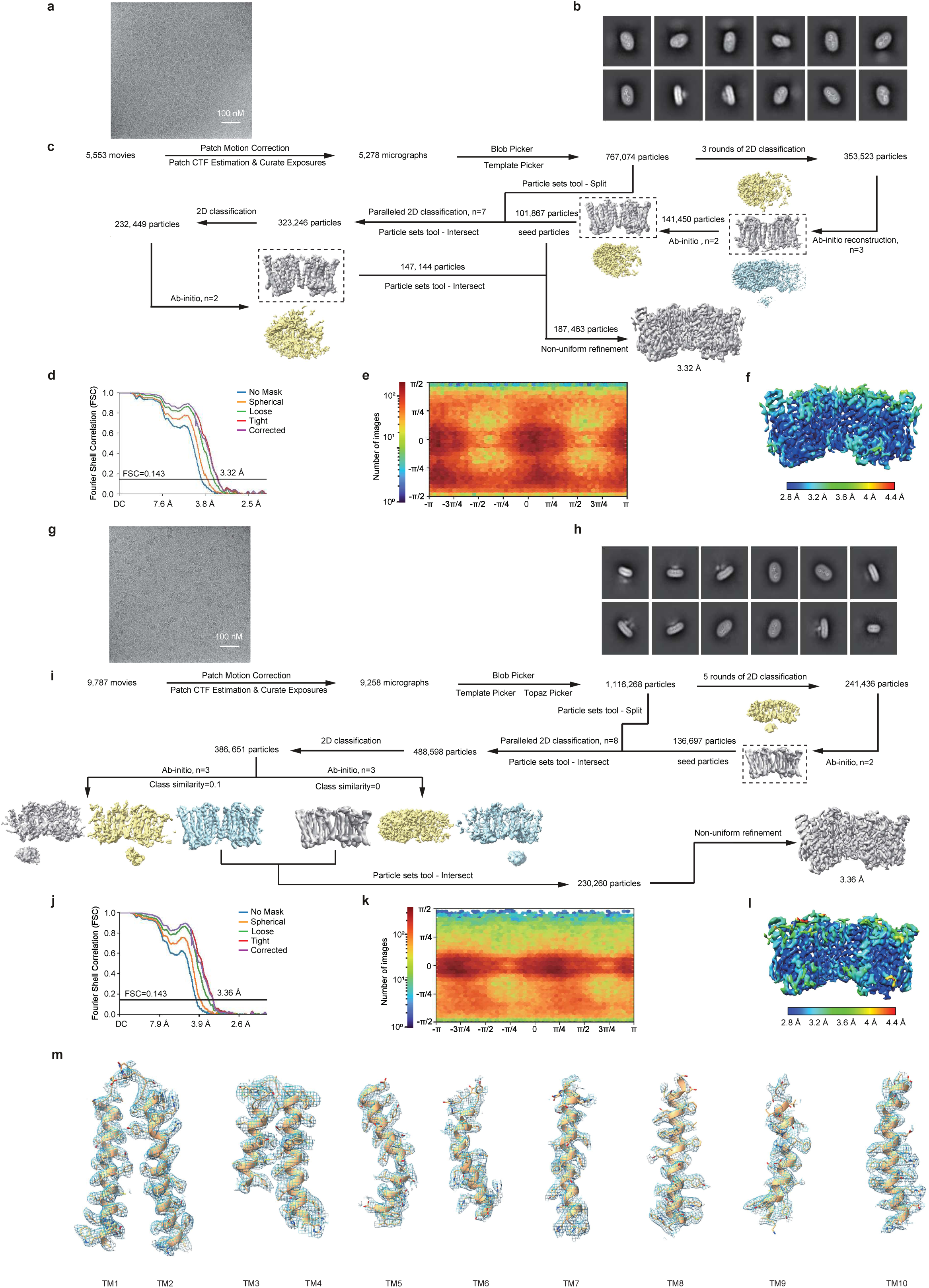
Data processing workflows of XPR1-KIDINS220 (1-432) in the closed state in the absence of KH_2_PO_4_ (XPR1^C1^) and the presence of 10 mM KH_2_PO_4_ (XPR1^C2^) **a**, A representative raw micrograph of XPR1-KIDINS220 (1-432) in the closed state in the absence of KH_2_PO_4_. **b**, Representative 2D class averages of the XPR1^C1^ state structure. **c**, Data processing pipeline for the XPR1^C1^ state structure. **d**, Gold-standard FSC curves of the XPR1^C1^ state structure. Resolution estimations were based on the criterion of the FSC 0.143 cutoff. **e**, Angular distribution of the final reconstruction of the XPR1^C1^ state structure. **f**, Local-resolution estimation of the final density map of the XPR1^C1^ state structure. **g**, A representative raw micrograph of XPR1-KIDINS220 (1-432) in the closed state in the presence of 10 mM KH_2_PO_4_. **h**, Representative 2D class averages of the XPR1^C2^ state structure. **i**, Data processing pipeline for the XPR1^C2^ state structure. **j**, Gold-standard FSC curves of the XPR1^C2^ state structure. Resolution estimations were based on the criterion of the FSC 0.143 cutoff. **k**, Angular distribution of the final reconstruction of the XPR1^C2^ state structure. **l**, Local-resolution estimation of the final density map of the XPR1^C2^ state structure. **m**, Representative cryo-EM densities of the transmembrane helices in the XPR1^C1^ state structure which are shown as mesh, superimposed with the corresponding stick models.

**Supplementary Fig. 3.**
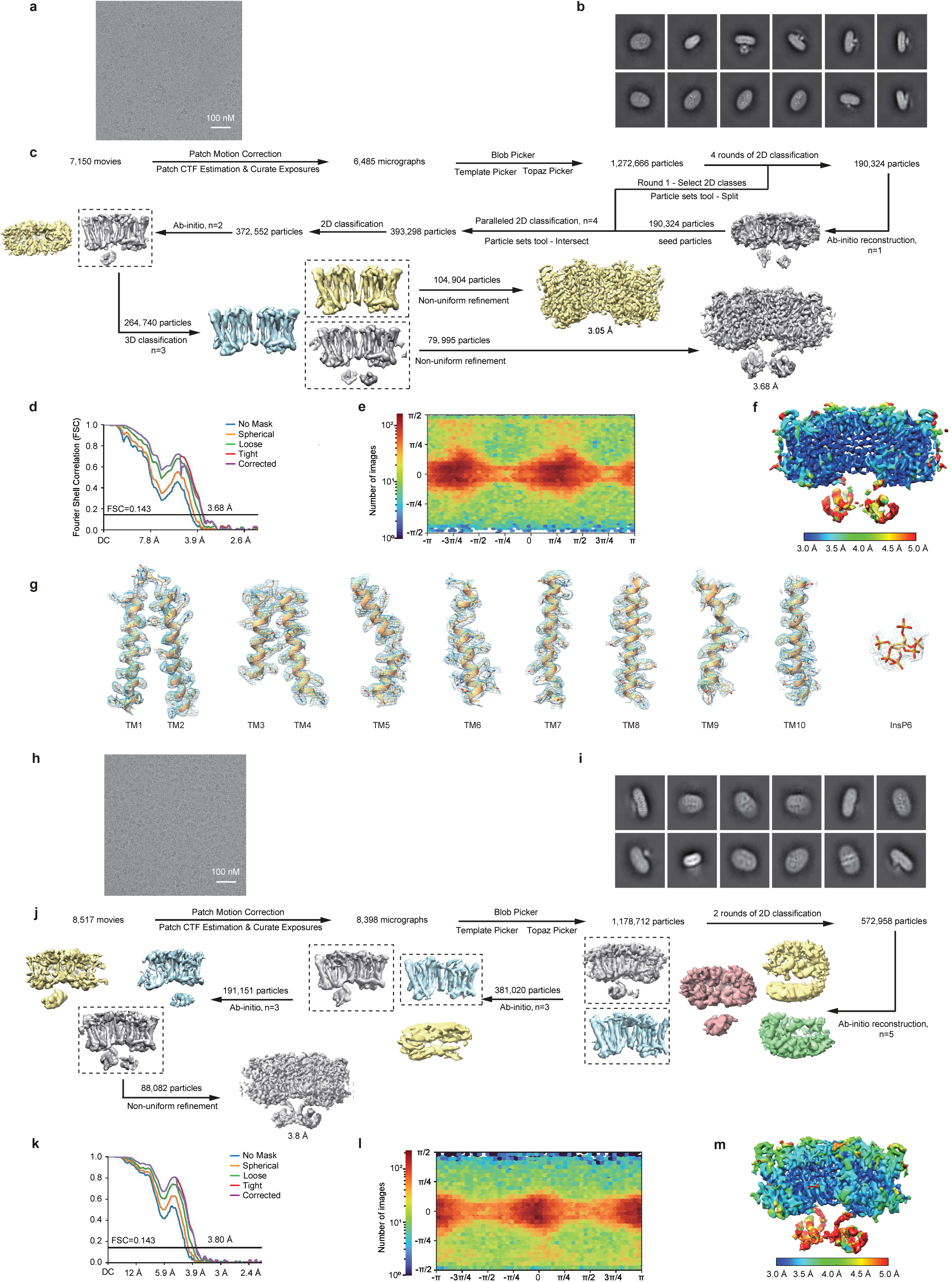
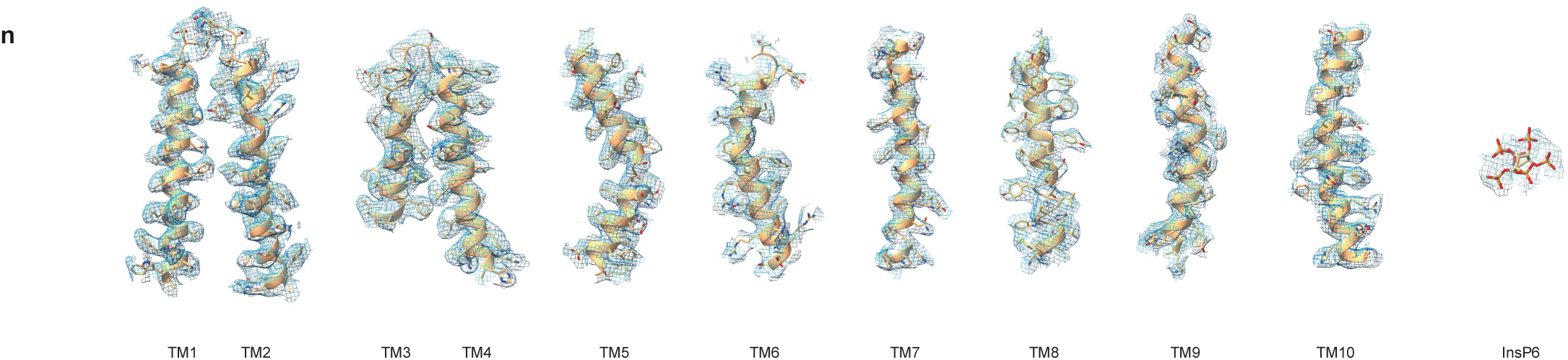
Data processing workflows of XPR1-KIDINS220 (1-432) in the outward-open state in the presence of 10 mM InsP6 and 10 mM KH_2_PO_4_ (XPR1^OUT^) and XPR1-KIDINS220 (1-432) in the closed state in the presence of 10 mM InsP6 and absence of KH_2_PO_4_ (XPR1^C_SPX^). **a**, A representative raw micrograph of XPR1-KIDINS220 (1-432) in the outward-open state in the presence of 10 mM InsP6 and 10 mM KH_2_PO_4_. **b**, Representative 2D class averages of the XPR1^OUT^ state structure. **c**, Data processing pipeline for the XPR1^OUT^ state structure. **d**, Gold-standard FSC curves of the XPR1^OUT^ state structure. Resolution estimations were based on the criterion of the FSC 0.143 cutoff. **e**, Angular distribution of the final reconstruction of the XPR1^OUT^ state structure. **f**, Local-resolution estimation of the final density map of the XPR1^OUT^ state structure. **g**, Representative cryo-EM densities of the transmembrane helices and InsP6 in the XPR1^OUT^ state structure which are shown as mesh, superimposed with the corresponding stick models. **h**, A representative raw micrograph of XPR1-KIDINS220 (1-432) in the closed state in the presence of 10 mM InsP6 and absence of KH_2_PO_4_. **i**, Representative 2D class averages of the XPR1^C_SPX^ state structure. **j**, Data processing pipeline for the XPR1^C_SPX^ state structure. **k**, Gold-standard FSC curves of the XPR1^C_SPX^ state structure. Resolution estimations were based on the criterion of the FSC 0.143 cutoff. **l**, Angular distribution of the final reconstruction of the XPR1^C_SPX^ state structure. **m**, Local-resolution estimation of the final density map of the XPR1^C_SPX^ state structure. **n**, Representative cryo-EM densities of the transmembrane helices and InsP6 in the XPR1^C_SPX^ state structure which are shown as mesh, superimposed with the corresponding stick models.

**Supplementary Fig. 4.**
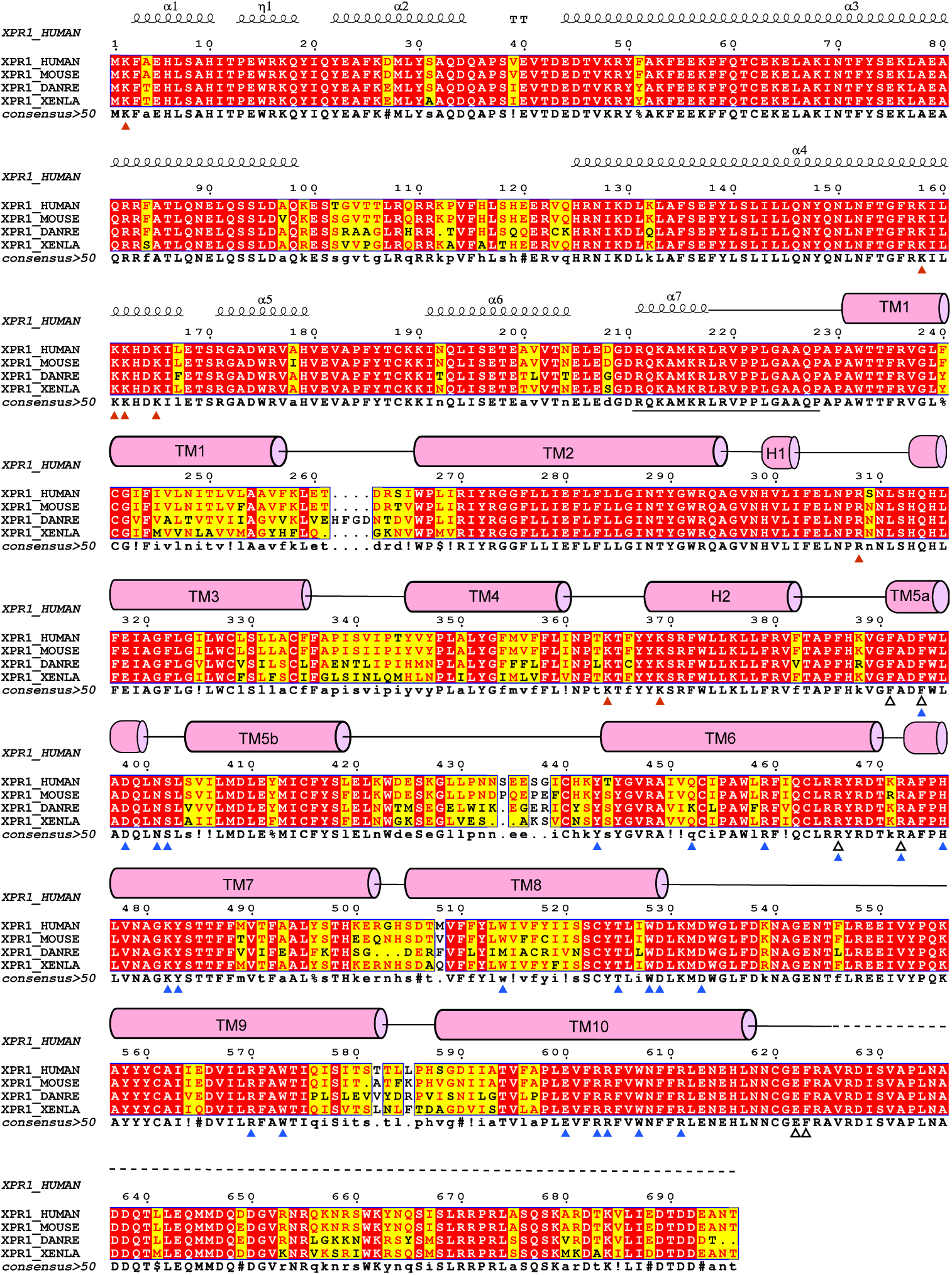
Sequence alignment of the XPR1 protein homologs among multiple species. Sequence alignment of the residues of XPR1 protein homologs among human (*Homo sapiens*), mouse (*Mus terricolor*), zebrafish (*Danio rerio*) and African clawed frog (*Xenopus laevis*). Residues related with the InsP6 binding pocket, the phosphate ion transport pore and the C-terminal plug-in loop were indicated with red, blue, and hollow triangles under the sequence, respectively.

**Supplementary Fig. 5.**
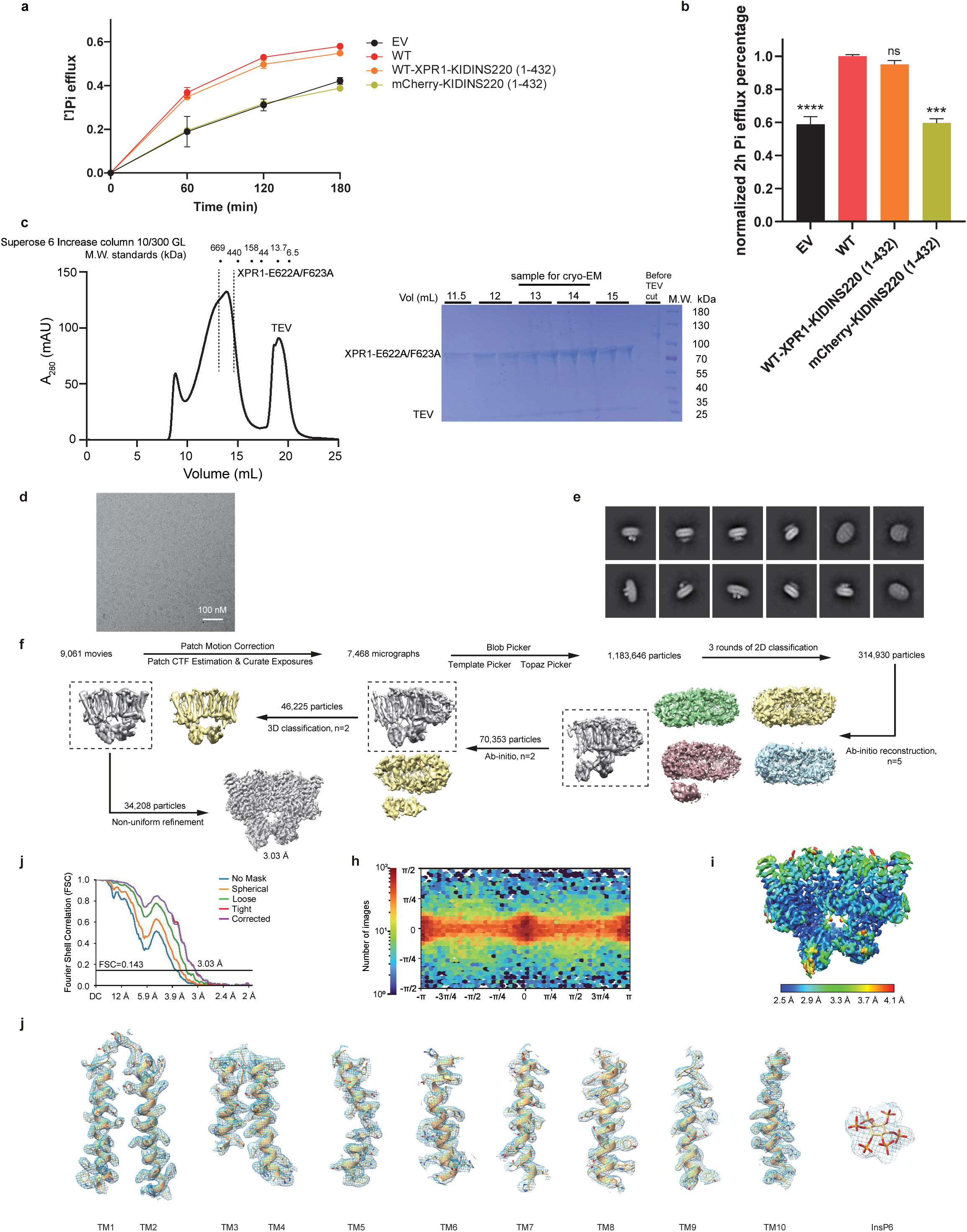
The effect of KIDINS220 (1-432) on XPR1’s Pi export activity and data processing workflows of XPR1-E622A/F623A mutant in the inward-facing state in the presence of 10 mM InsP6 and 10 mM KH_2_PO_4_ (XPR1^In^) **a**, ^32^Pi efflux of HEK293T cells transfected with empty vector and constructs of WT-XPR1, WT-XPR1-KIDINS220 (1-432) and mCherry-KIDINS220 (1-432). Samples were taken and measured at indicated time points and the ratios of ^32^Pi against the total cellular ^32^Pi were calculated. Data are shown as mean ± s.d.; n≥3 biologically independent experiments. **b**, ^32^P efflux percentages of EV, WT-XPR1-KIDINS220 (1-432) and mCherry-KIDINS220 (1-432) in **a** at 2 h normalized against the values of WT-XPR1 obtained under identical conditions from the same experimental batch. Data are shown as mean ± s.d.; n≥3 biologically independent experiments. *P* values (ns ≥ 0.05, \**P* < 0.05, \*\**P* 0.01, \*\*\**P*< 1 × 10^−3^, \*\*\*\**P*< 1 × 10^−4^) were obtained by unpaired *t* test with Welch’s correction between EV, WT-XPR1-KIDINS220 (1-432), mCherry-KIDINS220 (1-432) and WT-XPR1. **c**, Size-exclusion chromatography profile and SDS-PAGE gel of purified XPR1-E622A/F623A mutant in the inward-facing state in the presence of 10 mM InsP6 and 10 mM KH_2_PO_4_. Fraction within dashed lines were concentrated for cryo-EM sample preparation. The molecular weight standards of the Superose 6 Increase column 10/300 GL were labeled on the SEC profile. **d**, A representative raw micrograph of the XPR1^In^ state structure. **e**, Representative 2D class averages of the XPR1^In^ state structure. **f**, Data processing pipeline for the XPR1^In^ state structure. **g**, Gold-standard FSC curves of the XPR1^In^ state structure. Resolution estimations were based on the criterion of the FSC 0.143 cutoff. **h**, Angular distribution of the final reconstruction of the XPR1^In^ state structure. **i**, Local-resolution estimation of the final density map of the XPR1^In^ state structure. **j**, Representative cryo-EM densities of the transmembrane helices and InsP6 in the XPR1^In^ state structure which are shown as mesh, superimposed with the corresponding stick models.

**Supplementary Fig. 6.**
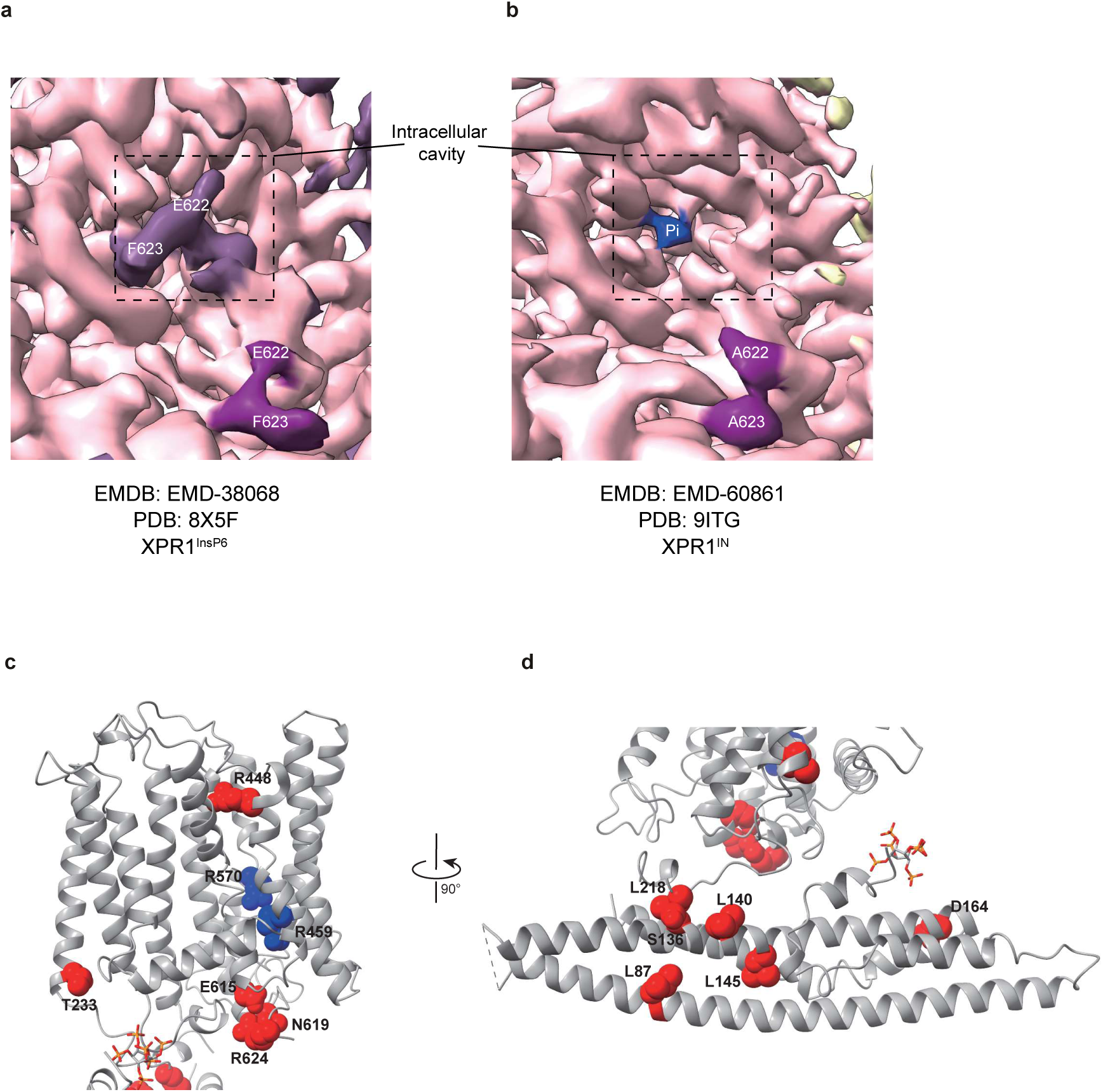
The cryo-EM densities around the intracellular cavities and the reported pathogenic mutations of XPR1 in PFBC patients. **a**, The density map around the intracellular cavity of XPR1^InsP6^, as reported by Jiang et al.^34^. The density of the Glu622/Phe623 motif in their atomic model was colored in magenta. The density in the intracellular cavity which hadn’t been built with atomic model was colored in purple. The Glu622/Phe623 residues were labeled on the map. **b**, The density map around the intracellular cavity of XPR1^In^ in our study. The densities of Ala622/Ala623 (E622A/F623A) and the phosphate ion in our atomic models were colored in magenta and blue, respectively. **c**,**d**, The single amino acid substitutions in the TMD (**c**) or the SPX domain (**d**) of XPR1 reported in PFBC patients. The residues involved in this study were presented as blue spheres and the other mutations as red spheres. The substitutions found in patients are: L87P, S136N, L140P, L145P, D164Y, L218S, T233S, D262Efs*6-R448W, R459C, R570C, R570L, E615K, N619D and R624H.

**Supplementary Table 1.** Cryo-EM data collection, refinement and validation statistics.

| PDB ID<br>EMDB ID | XPR1<sup>C1</sup><br>9INE<br>EMD-60704 | XPR1<sup>C2</sup><br>9INF<br>EMD-60705 | XPR1<sup>OUT</sup><br>9INH<br>EMD-60707 | XPR1<sup>C_SPX</sup><br>9IUC<br>EMD-60897 | XPR1<sup>IN</sup><br>9ITG<br>EMD-60861 |
| --- | --- | --- | --- | --- | --- |
| **Data collection and processing** |  |  |  |  |  |
| Magnification | 165,000 × | 165,000 × | 96,000 × | 165,000 × | 165,000 × |
| Voltage (kV) | 300 | 300 | 300 | 200 | 300 |
| Electron exposure (e<sup>-</sup>/Å<sup>2</sup>) | 50 | 50 | 50 | 50 | 50 |
| Defocus range (μm) | -1.5 to -1.8 | -1.5 to -1.8 | -1.5 to -1.8 | -1.5 to -1.8 | -1.5 to -1.8 |
| Pixel size (Å) | 0.821 | 0.821 | 0.808 | 0.74 | 0.74 |
| Symmetry imposed | C2 | C2 | C2 | C2 | C2 |
| Initial particle images (no.) | 767,074 | 1,116,268 | 1,272,666 | 1,178,712 | 1,183,646 |
| Final particle images (no.) | 187,463 | 230,260 | 79,995 | 88,082 | 34,208 |
| Map resolution (Å) | 3.32 | 3.36 | 3.68 | 3.8 | 3.03 |
| FSC threshold | 0.143 | 0.143 | 0.143 | 0.143 | 0.143 |
| Map resolution range (Å) | 1.751-32.947 | 1.805-38.583 | 2.101-47.126 | 2.222-44.618 | 1.730-37.066 |
| **Refinement** |  |  |  |  |  |
| Initial model used (PDB code) | AF-Q9UBH6-F1 | 9INE | AF-Q9UBH6-F1 | 9INH | 9INH |
| Model resolution (Å) | 3.4 | 3.5 | 3.8 | 3.9 | 3.2 |
| FSC threshold | 0.5 | 0.5 | 0.5 | 0.5 | 0.5 |
| Model resolution range (Å) | 50-3.3 | 50-3.4 | 50-3.7 | 50-3.8 | 50-3.0 |
| Map sharpening *B* factor (Å<sup>2</sup>) | -134.9 | -142.6 | -116.7 | -141.1 | -81.8 |
| Model composition |  |  |  |  |  |
| Non-hydrogen atoms | 7520 | 7282 | 10982 | 10360 | 10435 |
| Protein residues | 802 | 802 | 1221 | 1140 | 1207 |
| Ligands | 26 | 18 | 34 | 26 | 16 |
| *B* factors (Å<sup>2</sup>) |  |  |  |  |  |
| Protein | 70.06 | 75.01 | 60.56 | 126.90 | 79.75 |
| Ligand | 72.49 | 73.75 | 54.15 | 102.55 | 78.32 |
| R.m.s. deviations |  |  |  |  |  |
| Bond lengths (Å) | 0.004 | 0.003 | 0.004 | 0.006 | 0.014 |
| Bond angles (°) | 0.623 | 0.672 | 0.698 | 0.958 | 1.648 |
| Validation |  |  |  |  |  |
| MolProbity score | 1.96 | 1.90 | 1.77 | 1.81 | 2.36 |
| Clash score | 14.56 | 14.57 | 11.33 | 12.68 | 12.23 |
| Poor rotamers (%) | 0.56 | 0.56 | 1.01 | 0.10 | 3.57 |
| Ramachandran plot |  |  |  |  |  |
| Favored (%) | 95.74 | 96.49 | 96.86 | 96.79 | 95.12 |
| Allowed (%) | 4.26 | 3.51 | 3.14 | 3.21 | 4.88 |
| Disallowed (%) | 0 | 0 | 0 | 0 | 0 |

**Supplementary Table 2.** Sample conditions of the cryo-EM structures in our study.

| PDB ID<br>EMDB ID | XPR1<sup>C1</sup><br>9INE<br>EMD-60704 | XPR1<sup>C2</sup><br>9INF<br>EMD-60705 | XPR1<sup>OUT</sup><br>9INH<br>EMD-60707 | XPR1<sup>C_SPX</sup><br>9IUC<br>EMD-60897 | XPR1<sup>IN</sup><br>9ITG<br>EMD-60861 |
| --- | --- | --- | --- | --- | --- |
| Wild-type XPR1 | + | + | + | + | - |
| XPR1-E622A/F623A mutant | - | - | - | - | + |
| KIDINS220 (1-432) | + | + | + | + | - |
| 10 mM KH<sub>2</sub>PO<sub>4</sub> | - | + | + | - | + |
| 10 mM InsP6 | - | - | + | + | + |

